# Primate-specific adaptation of Ku protects transcriptomic integrity by suppressing Alu-mediated alternative splicing

**DOI:** 10.64898/2025.12.17.694518

**Authors:** Tianji Yu, Jimin Yoon, Yimeng Zhu, Angelina Li, Brian J. Lee, Daniel F. Moakley, Jing Duan, Qiannian Deng, Fanjia Hou, Mengfang Yan, Vincenzo A. Gennarino, Ke Zhang, Li Chen, Shan Zha, Chaolin Zhang

## Abstract

Accurate pre-mRNA splicing is essential for the transfer of genetic information but faces unique challenge in higher primates due to the massive expansion of intronic Alu elements^1–3^. While studying Ku, the Ku70/Ku80 heterodimer best known for initiating non-homologous end-joining (NHEJ) by encircling DNA ends^4^, we discovered that Ku expression increased markedly during primate evolution in parallel with Alu expansion^5^. Ku binds double-stranded RNA (dsRNA) stem-loops, including those at the antisense Alu (asAlu) elements within introns^5^. Here, we show that Ku-depletion in human cells has a broad impact on splicing largely independent of cell-cycle states, NHEJ, or innate immune signaling, significantly affecting ~8-10% of quantifiable alternative splicing events. Mechanistically, Ku directly binds exonic asAlu to prevent their aberrant inclusion and binds asAlu within inverted-repeat Alu (irAlu) pairs flanking canonical exons to prevent exon skipping^6^. Among human tissues, Ku expression in the brain is consistently ~50% lower, correlating with more permissive expression of Alu-derived splice variants, particularly those encoding mitochondrial proteins and RNA-binding factors. Correspondingly, heterozygous Ku loss in patient causes developmental delay, neurological dysfunction, and acidosis. Together our findings identified Ku as a critical suppressor of Alu-associated alternative splicing co-opted during evolution with implications for primate brain function and human disease.

---

Pre-mRNA splicing is tightly regulated by RNA-binding proteins (RBPs) and is essential for the accurate and efficient flow of genetic information from DNA to protein^7–10^. At the same time, by creating, removing, and reshuffling exons, the widespread and divergent alternative splicing in mammals expands transcriptomic diversity and contributes to mammalian evolution^11–13^. Most new transcripts arising from alternative splicing are likely evolutionarily neutral or deleterious and are eliminated by natural selection, while a subset acquires adaptive fitness^14,15^. Higher primates face a unique challenge to ensure accurate splicing due to the massive expansion of Alu elements in gene-rich regions^1,2^. Alu repeats constitute ~11% of the human genome (~1.4 million copies) and are enriched in promoters, introns, and 3’ untranslated regions (UTRs). Each ~300-bp Alu element contains two conserved stem–loops (SLs) linked together and followed by a polyadenine (polyA) tract. Depending on its orientation relative to the host gene, an Alu can be transcribed in either sense (sAlu) or antisense (asAlu) direction. Because asAlu sequences frequently harbor cryptic splice sites as well as polyuridine (polyU) tracts, if not properly regulated, they are prone to “exonization,” leading to inappropriate inclusion of Alu-derived sequences in mature mRNA^3,16,17^. Moreover, two intronic Alu elements in the opposite orientations can anneal with each other to form inverted-repeat Alu (irAlu), a long double-stranded RNA (dsRNA) duplex that can be immunogenic by activating MDA5^18,19^. When such irAlu-mediated dsRNA spans a pre-existing exon, the intervening exon can be looped out, resulting in “aberrant” exon skipping^20–22^. In the case of the *TBXT* gene encoding a T-box transcription factor that is critical for embryonic development, irAlu-mediated skipping of exon 6 in hominoids has been shown to underlie the evolutionary tail loss^22^. Not surprisingly, multiple mechanisms safeguard against aberrant splicing associated with Alu expansion in the pre-mRNAs. For example, hnRNP C binds to the polyU tracks within asAlu and competes with U2AF65 to repress the inclusion of asAlu-derived exons^23^. In addition, RNA helicase DHX9 unwinds the irAlu-duplex to prevent circular RNA formation and aberrant transcriptome rewiring^24^. However, both hnRNP C and DHX9 are themselves highly conserved in mammals. It remains unknown whether a primate-specific mechanism co-opted with Alu expansion to cope with the widespread RNA secondary structures, both within individual Alu stem–loops and between paired irAlu elements and thereby preserve RNA-processing fidelity and transcriptome integrity in primates, including humans.

While investigating the non-homologous end-joining (NHEJ) DNA double-strand break repair pathway, we uncovered a primate-specific functional exaptation of Ku, a heterodimeric ring-shaped complex of Ku70 and Ku80, as a nuclear dsRNA-binding protein (dsRBP)^5^. Canonically, Ku initiates NHEJ by encircling DNA ends and recruiting downstream factors such as DNA-PKcs and LIG4^4^. Paradoxically, despite the comparable genome sizes, human and higher primate cells express ~100-fold more Ku but not LIG4 or XRCC4, than lower primates predating Alu expansion (*e.g.*, lemur) or non-primate mammals *(e.g*., mouse and whale)^5,25,26^. Moreover, Ku is dispensable for murine development yet essential in >1,000 tested human cell lines^5,27–30^. Inducible degradation of Ku, but not LIG4, in human cells activates interferon (IFN) and NF-κB signaling via dsRNA sensors MDA5/RIG-I - MAVS^5^, followed by PKR-mediated translation suppression and OAS/RNase L-mediated RNA decay, ultimately contributing to growth arrest and cell death^5^. Protein-RNA interaction profiling using irCLIP revealed widespread Ku binding at human nuclear dsRNA stem-loops derived from snoRNA, rRNA, tRNA, as well as intronic and 3’UTR regions of protein-coding genes^5,31^, with a striking enrichment at the two conserved stem loops of primate-specific Alu elements, almost exclusively in the antisense orientation (98%)^5^. Concordantly, the increased expression of *XRCC6* and *XRCC5,* which encodes Ku70 and Ku80, tightly correlates with Alu expansion (r = 0.94/0.95)^5^. These findings identify Ku as a nuclear dsRBP that is upregulated in higher primates to accommodate Alu expansion in nuclear RNA. Yet the consequences of Ku binding for RNA processing in human and primate cells remain unknown. Because Ku forms a closed ring that encircles dsRNA, we reasoned that its binding could stabilize stem-loops and thereby modulate RNA processing events that rely on dynamic pairing and unpairing between local or distal sequences. Given that >65% of Ku-bound sites lie in introns^5^, including Alu elements widely implicated in primate-specific alternative splicing^1–3,5^, and that Ku proteomic interactomes in human cells are enriched for RNA-binding proteins, including splicing factors^32,33^, here we investigate how Ku co-option with Alu expansion regulates pre-mRNA splicing to safeguard transcriptome integrity while allowing potential molecular innovations during primate evolution.

## Result

### Ku is a major splicing regulator influencing ~8-10% of alternative splicing events in human cells

To investigate the role of Ku in pre-mRNA splicing, we used previously characterized HCT116 clones in which endogenous Ku80 is tagged with an auxin-inducible degron (AID) for rapid and inducible degradation^5^. Because the N-terminal AID tag and selection cassette separate the upstream transcription start site from a downstream transcriptional enhancer element, these cells grow normally but express Ku protein at ~25% of wild-type levels before auxin (IAA) induction, providing an opportunity to assess the dose-dependent impact of Ku on RNA processing (Fig. 1a). AID induction with IAA and Dox reduces Ku80 protein to ~5% within 24 hours and ~1% in 4 days (Fig. 1b, Extended Data Fig. 1a). To allow newly synthesized mRNAs upon Ku depletion to replace pre-existing transcripts, we performed our initial analyses by deep RNA-sequencing (RNA-seq) using RNA collected four days after AID induction, when cellular viability remained adequate^5^ (Extended Data Table 1).

**Figure 1:**
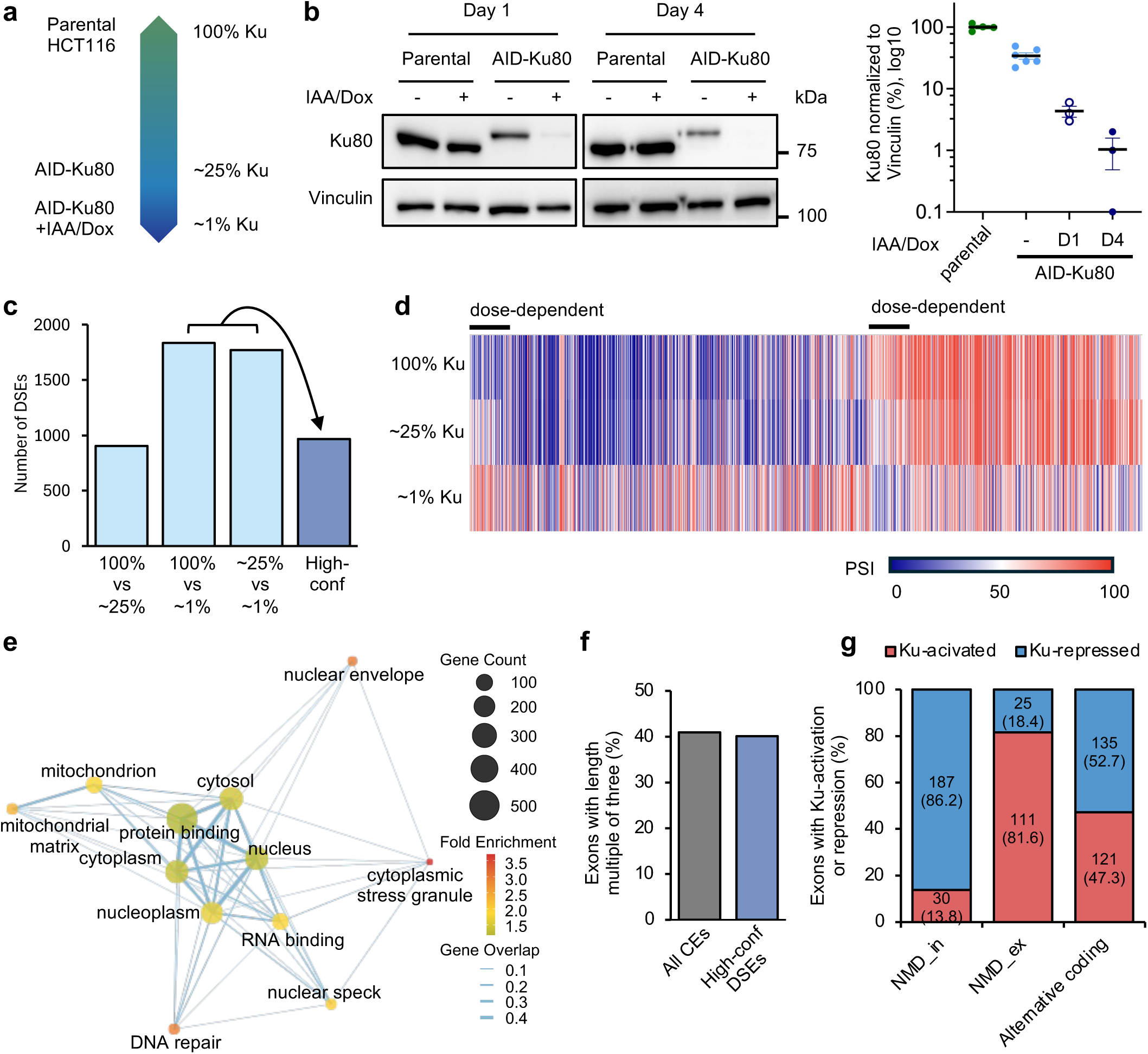
Ku depletion results in global splicing changes in human cells in a dose-dependent manner. Only cassette exons (CE) are shown in this figure. **a,** Ku80 protein levels under different conditions. Compared with parental HCT116 cells (100% Ku), Ku80 protein level reduces to ~25% in AID-Ku80-expressing HCT116 cells without Dox and IAA treatment and to ~1% with Dox and IAA treatment at day 4. **b,** Left: Representative western blot analyses of Ku80 protein in parental and AID-Ku80-expressing HCT116 cells at day 1 and 4, with or without IAA & Dox treatment to induce Ku depletion. Right: Quantification of Ku80 protein levels normalized to Vinculin from independent Western blot replicates. Y-axis is displayed on a log scale. **c,** Differentially spliced events (DSEs) detected upon different levels of Ku80 (100% vs. ~25%, 100% vs. ~ 1%, and ~25% vs. ~1%; light blue bars). DSEs shared between 100% vs. ~1% Ku and ~25% vs. ~1% Ku comparisons with consistent direction of change are defined as high-confident DSEs (dark blue bar). **d,** Heatmap showing the PSI values of 967 high-confidence DSEs. A subset of exons showing dose-dependent changes between 100% vs. ~25% Ku are indicated with bars at the top. **e,** Gene ontology (GO) terms enriched in genes containing high-confidence DSEs. Each node represents a GO term. Each edge connecting two GO terms represents fraction of overlapping genes. **f,** The percentage of exons with lengths that are multiple of three among all cassette exons (control) or high-confidence DSEs. **g**, Number of Ku-repressed (blue) and Ku-activated (red) high-confident DSEs grouped by NMD impact. Only exons supported by full-length transcripts with open reading frame annotations were included in this analysis. NMD_in: inclusion of the alternative exon causes NMD. NMD_ex: exclusion of the alternative exon causes NMD. Alternative coding: alternative splicing generates two coding transcripts.

Differential splicing analysis with the Quantas pipeline^34,35^ detected six major classes of alternative splicing events: cassette exons, tandem cassette exons, alternative 5’ or 3’ splice-sites, mutually exclusive exons, and retained introns using stringent cut-offs (changes in percent spliced-in (|ΔPSI|) ≥ 10, FDR ≤ 0.05, and exon-junction read coverage ≥ 20). Across three independently derived AID-Ku80 clones, Ku degradation (from ~25% to ~1% of normal levels) resulted in 3,605 differential splicing events (DSEs), representing 7.9% of all quantifiable events. A comparable number of DSEs was observed when using the parental HCT116 cells with full Ku expression as control (100% Ku vs. ~1% Ku; 3,649 DSEs or 8% of all quantifiable events) (Extended Data Fig. 1b and Extended Data Table 2). Among different types of alternative splicing events, cassette exon inclusion/skipping events are the most abundant, accounting for around half of all DSEs, with 1,772 significant events in the isogenic clones upon inducible Ku-degradation (25% vs. ~1% Ku) and 1,833 events when compared to parental AID control (100% vs. ~1% Ku). These changes represent approximately 10% of all quantifiable cassette exons in HCT116 cells (Fig. 1c, Extended Data Tables 3 and 4). Consistent with a dose-dependent effect, loss of ~75% of normal Ku levels (100% vs. ~25%) affected a smaller number (1,886) of DSEs (vs.36,49 DSEs when losing ~99% Ku), including 902 cassette exons (Fig. 1c and Extended Data Fig. 1b). To confirm these findings, we used an independently derived HCT116 cell line with conditional Ku80 deletion via the Cre-LoxP system (*Ku86^flox/-^:CreER-T2* and control *Ku86^flox/-^*)^5,36^ (Extended Data Fig. 1c). Comparison of cells with heterozygous vs homozygous Ku knockout (50% vs 0% of normal Ku) identified 2,854 total DSEs, including 1,352 cassette exons (vs. 1,833 cassette exons between 100% vs. ~1% Ku, Extended Data Fig. 1d), supporting a robust and consistent role of Ku in regulating RNA splicing in human cells.

To define the mechanism by which Ku regulates alternative splicing and their biological impacts on transcriptome integrity, we focused on cassette exons. Overlapping the significant events from the 100% vs. ~1% Ku and ~25% vs. ~1% Ku comparisons in the AID-Ku system identified 968 exons; all of them, except for one, showed consistent directions of splicing changes, resulting in 967 high-confidence Ku-dependent cassette exons used for further analysis (Fig. 1c). Of these, 577 exons (~60%) were repressed by Ku (PSI increased upon Ku depletion), whereas 390 (~40%) were activated by Ku (PSI decreased upon Ku depletion) (Extended Data Fig. 1e). Most of the high-confidence Ku-dependent exons defined here had comparable splicing differences in magnitude between the 100% vs. ~1% Ku and between ~25% vs. ~1% Ku), whereas 91 exons (~10%) showed significant differential splicing between 100% vs. ~25% Ku conditions, reflecting dose-dependent splicing regulation (Fig. 1d and Extended Data Fig. 1e,f). The 967 high-confidence Ku-dependent exons are from 710 unique genes. Gene ontology (GO) analysis revealed a significant enrichment of mitochondrial genes (133 exons in 99 genes, 13.9%, Benjamini FDR<1.3e-6; e.g., *COQ3* and *MTO1*), RNA-binding proteins (49 exons in 36 genes, 5.1%, Benjamini FDR<2.6e-7; e.g., *MBNL2*, *EXOSC9* and *UA2F1L4*), and DNA repair factors (38 exons from 29 genes, 4.1%, Benjamini FDR<1.8e-4; e.g., *POLB* and *RAD1*) (Fig. 1e; see examples in Extended Data Fig. 2a-d, and gene lists in Extended Data Table 5), indicating a broad functional impact of Ku on multiple downstream molecular pathways.

The inclusion of Ku-repressed exons or the skipping of Ku-activated exons can be either in-frame, altering protein structure, or out-of-frame, reducing gene expression by introducing frameshifts that cause premature termination and trigger nonsense-mediated decay (NMD)^37^. In this context, ~40% of Ku-dependent exons are predicted to preserve the reading frame (i.e., exon lengths in multiples of three), whereas the remaining ~60% are expected to introduce frameshifts (Fig. 1f). Consistently, analysis of premature-termination–codon (PTC)-containing transcripts subject to NMD revealed that 58% of high-confidence Ku-dependent exons indeed generate NMD-targeted transcripts upon Ku-deletion through either aberrant exon inclusion (NMD_in) or exon exclusion (NMD_ex). Notably, in 84.4% of these NMD-coupled cassette exons (Ku-repressed NMD_in and Ku-activated NMD_ex exons), Ku suppresses the unproductive NMD isoforms (Fig. 1g), underscoring a key role for Ku in maintaining the integrity of the protein-coding transcriptome by ensuring highly efficient production of the canonical splice variants.

### Ku directly represses the inclusion of asAlu-derived alternative exons

Given Ku’s strong binding preference for asAlu elements^5^ and the established role of Alu in primate-specific alternative splicing^3,16,17,20–22^, we categorized the Ku-dependent cassette exons by the presence and location of Alu elements. Among the 967 high-confidence Ku-dependent exons, the frequencies of exons containing Alu in their flanking introns (intronic Alu, 742/967, 76.7%) or lacking Alu entirely (no Alu, 205/967, 21.2%) largely matched their abundances in the human genome, but exonic Alu-containing exons (165/967, 17.1%) showed a distinctive pattern. As expected from the presence of polyU and cryptic splice sites uniquely in asAlu, exonic sAlu exons were rare in human transcriptome (1%), whereas exonic asAlu exons account for 6.3% of all cassette exons (Fig. 2a). Strikingly, among the Ku-dependent exons, exonic asAlu accounts for 158/967 (16.3%), representing a 2.6-fold enrichment (p=1.07e-28; Fisher-exact test), while exonic sAlu accounts for 7/967 (0.72%) largely as expected (Fig. 2a). Consistently, among cassette exons, those with overlapping asAlu are more likely to be Ku-dependent, as compared to exons without Alu (do not have Alu in the exons or flanking introns) (odds ratio=4.5; Extended Data Fig. 3a) Moreover, most asAlu-derived exons (149 of 158, 94.3%) were repressed by Ku, exhibiting increased PSI upon Ku degradation, compared to 67.8% of non-Alu exons (p=1.02e-10, Fisher’s exact test) and ~56.6% of exons with Alu in their flanking introns (Fig. 2b,c). Among the nine exonic asAlu exons “oddly” activated by Ku (9/158, 5.7%), 6 contain intronic Alu elements that might complicate their overall splicing outcomes (see below). Since Ku binds to individual stem-loops within asAlu elements, our data suggest a model that Ku degradation exposes the cryptic splice sites and polyU tracts within asAlu, thereby enabling aberrant exon inclusion (Fig. 2d). Indeed, RNA-seq data and RT–PCR validated the dose-dependent increase in PSI of four Ku-dependent exonic asAlu exons—such as *EXOSC9*—following Ku depletion in both AID-Ku80 and *Ku86^flox/-^:Cre-ERT2* cells (Fig. 2e,f,i, Extended Data Fig. 4a). Specifically, Ku depletion caused a 51-nt (17 amino acids) in-frame insert (PSI from 0.22 to 0.77, 100% to <0.1% Ku) into an intrinsically disordered region of human *EXOSC9* to generate isoform 2 unique for humans and not found in other model systems^38^ (Fig. 2e and Extended Data Fig. 4b).

**Figure 2:**
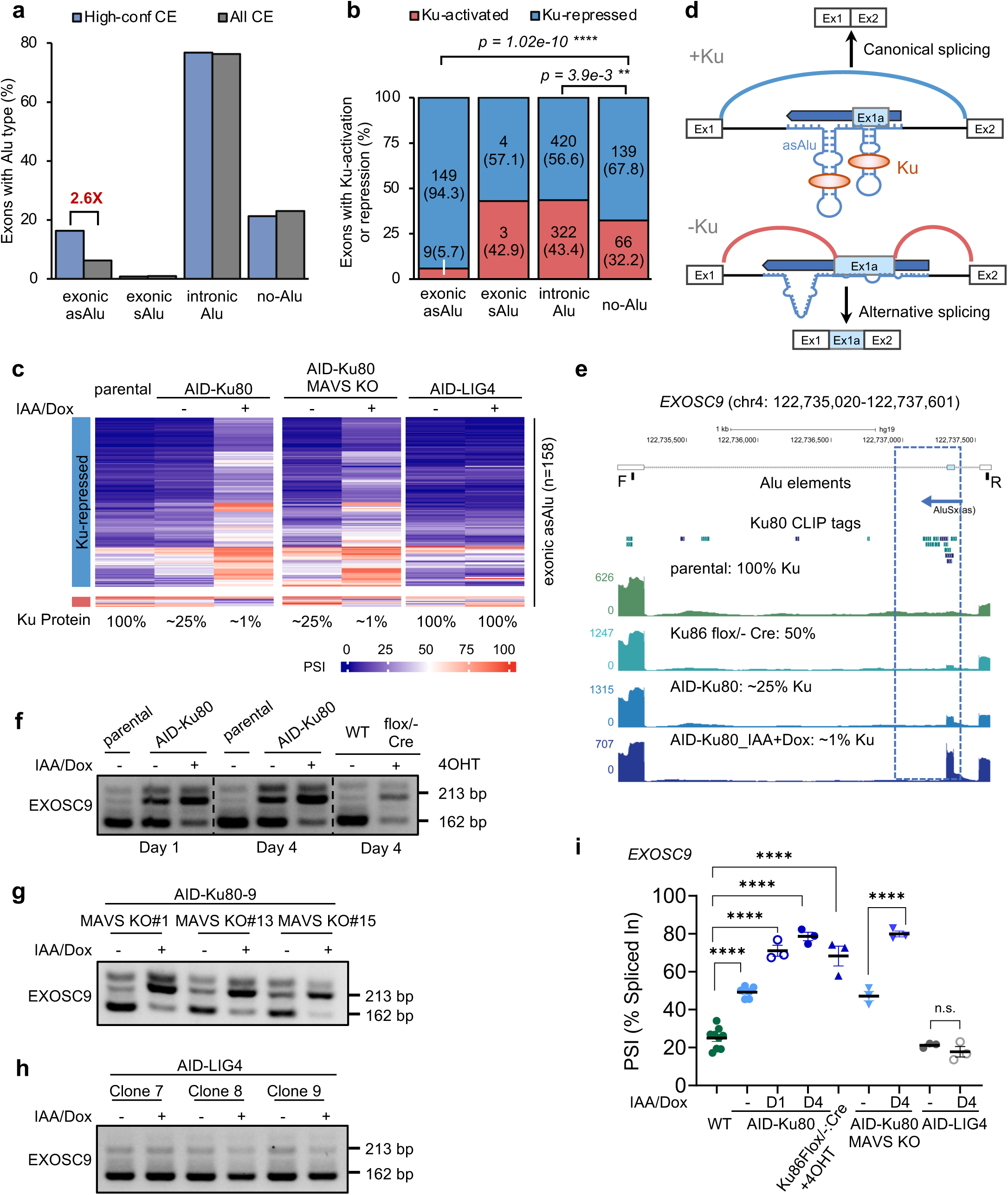
Ku directly represses inclusion of asAlu-derived alternative exons. **a,** Percentage of high-confident cassette exons containing exonic asAlu, exonic sAlu, intronic Alu, and no Alu in the exon or flanking introns (blue bars). All cassette exons (gray bars) are used for comparison. Note that a cassette exon can have both exonic and intronic Alu elements and can be assigned to multiple groups here. **b,** Percentage of Ku-repressed and Ku-activated high-confident DSEs grouped by type of Alu elements contained: exonic asAlu, exonic sAlu, intronic Alu (note cassette exons also containing exonic Alus were not excluded here) and no-Alu. For each Alu-type, the number and percentage (in parenthesis) of exons are also indicated. Statistics showing bias towards Ku-activated or Ku-repressed exons associated with specific Alu-types are also indicated (Fisher’s exact test p<0.0001 ****; p>0.05 ns). **c,** PSI values for high-confidence Ku-repressed (blue) and Ku-activated cassette exons (red) containing exonic asAlu in AID-Ku80 cells at 100%, ~25%, and ~1% Ku80 protein levels. PSI values in MAVS KO; AID-Ku80 cells without or with Ku depletion (~25% and ~1% Ku) and AID-LIG4 cells with or without LIG4 depletion are also shown. **d,** Schematic illustration of the proposed mechanism by which Ku regulates alternative splicing. Ku represses exon inclusion by directly binding to the stem loops of monomeric asAlu elements, thereby limiting splice-site accessibility and resulting in exon skipping. The exon inclusion level increases upon Ku depletion due to destabilization of the stem loops and increased accessibility of the splicing signals. **e,** UCSC Genome Browser view of the *EXOSC9* cassette exon containing an exonic asAlu element. Shown are the positions of the asAlu, Ku80 CLIP-seq binding sites, and RNA-seq read coverage in cells with varying Ku levels (100%, 50%, ~25%, and ~1%), illustrating the dosage-dependent impact of Ku depletion on exon inclusion. **f-h,** Representative RT-PCR validation data for *EXOSC9* transcripts in parental and AID-Ku80-expressing HCT116 cells on day 1 and day 4, with or without IAA and Dox treatment for Ku depletion, and *Ku86^flox/-^;CreERT2* HCT116 cells, with or without 4-hydroxy tamoxifen (4OHT) treatment for Ku depletion **(f)**, three clones of AID-Ku80-expressing MAVS knockout HCT116 cells, with or without IAA and Dox treatment for Ku depletion **(g);** and three clones of AID-Lig4-expressing HCT116 cells, with or without IAA and Dox treatment for Lig4 depletion **(h)**. **i,** RT-PCR quantification of exon inclusion as percent-spliced-in (PSI) values from three independent biological replicates. Statistical significance of differential splicing was determined by the unpaired Student’s *t*-test *p*<0.0001 ****; *p*>0.05 ns.

Mechanistically, multiple lines of evidence suggest that this is a direct consequence of Ku binding at exonic asAlu in splicing repression. First, Ku-irCLIP tag can be found within the affected exonic asAlu as shown for *EXOSC9* (Fig. 2e and Extended Data Fig. 2b). Second, the PSI increase of asAlu-derived exons inversely correlates with Ku abundance: exon inclusion is highest with ~1% Ku, intermediate at ~25%, and lowest at 100% (Fig. 2 c,e,f,i). Third, the effect can be observed within 24 hours upon Ku degradation and in G1-arrested cells after 48 hours of Ku degradation (Fig. 2e,f,i and Extended Data Fig. 5a-e), excluding secondary or indirect cell-cycle influence. Fourth, although Ku degradation activates dsRNA-induced, MAVS-dependent interferon (IFN) signaling in HCT116 cells, CRISPR knockout of MAVS abolishes IFN induction without affecting Ku-dependent splicing of asAlu-exons (Fig. 2c,g,i, Extended Data Fig. 6a,c) demonstrating that the splicing changes are not downstream of IFN signaling and innate immune activation. Fifth, this phenotype is NHEJ-independent, as inducible degradation of LIG4, a core NHEJ factor downstream of Ku, does not trigger inclusion of asAlu-derived exons (Fig. 2c,h,i, Extended Data Fig. 6b,c), ruling out an NHEJ effect.

Taken together, these results demonstrate that in human cells, Ku directly binds the stem-loops within exonic asAlu RNA, masking their cryptic splice signals from the spliceosome and preventing aberrant exon inclusion. This mechanism accounts for 149 of the 967 (~15%) high-confidence Ku-dependent cassette exons observed. Not surprisingly, these exons are higher-primate specific (after Alu expansion), with a majority of them introduced before the split of hominoids and old-world monkey, whereas a subset is hominoid-specific (Extended Data Fig. 7a).

### Ku represses exon skipping by inhibiting flanking intronic irAlu pairing and exon looping

We next analyzed the largest class of Ku-dependent cassette exons—those containing Alu elements in flanking introns (742/967; 76.7%) (Fig. 2b). Of these, 145 also have overlapped exonic Alu (including 141 with exonic asAlu), which we excluded given the strong Ku-dependent repression of exonic asAlu observed, leaving 597 exons containing only intronic Alu elements (61.7% of 967) (Fig. 3a). Unlike asAlu-derived exons, which predominantly undergo Ku-dependent skipping, intronic-Alu associated exons were repressed (285/597; 47.7%) or activated (312/597; 52.3%) by Ku at similar frequencies (Fig. 3b), suggesting that Ku-dependent splicing here likely involves more than one mechanism. Given that irAlu pairs are known to form long-range RNA secondary structures and regulate splicing, we further classified the intronic Alus based on the availability and location of irAlu pairs: those with irAlu pair(s) that flank the cassette exon (irAlu-inter, 312, 52.3%), which are the largest group, those with irAlu pair(s) within one intron (irAlu-intra, 163, 27.3%), and those with no irAlu pair (others, 122, 20.4%) (Fig. 3a). We prioritized irAlu-inter over irAlu-intra, as such if an exon is flanked by irAlu-inter, regardless of whether it also contains irAlu-intra, it will be classified as irAlu-inter in the initial analyses.

**Figure 3:**
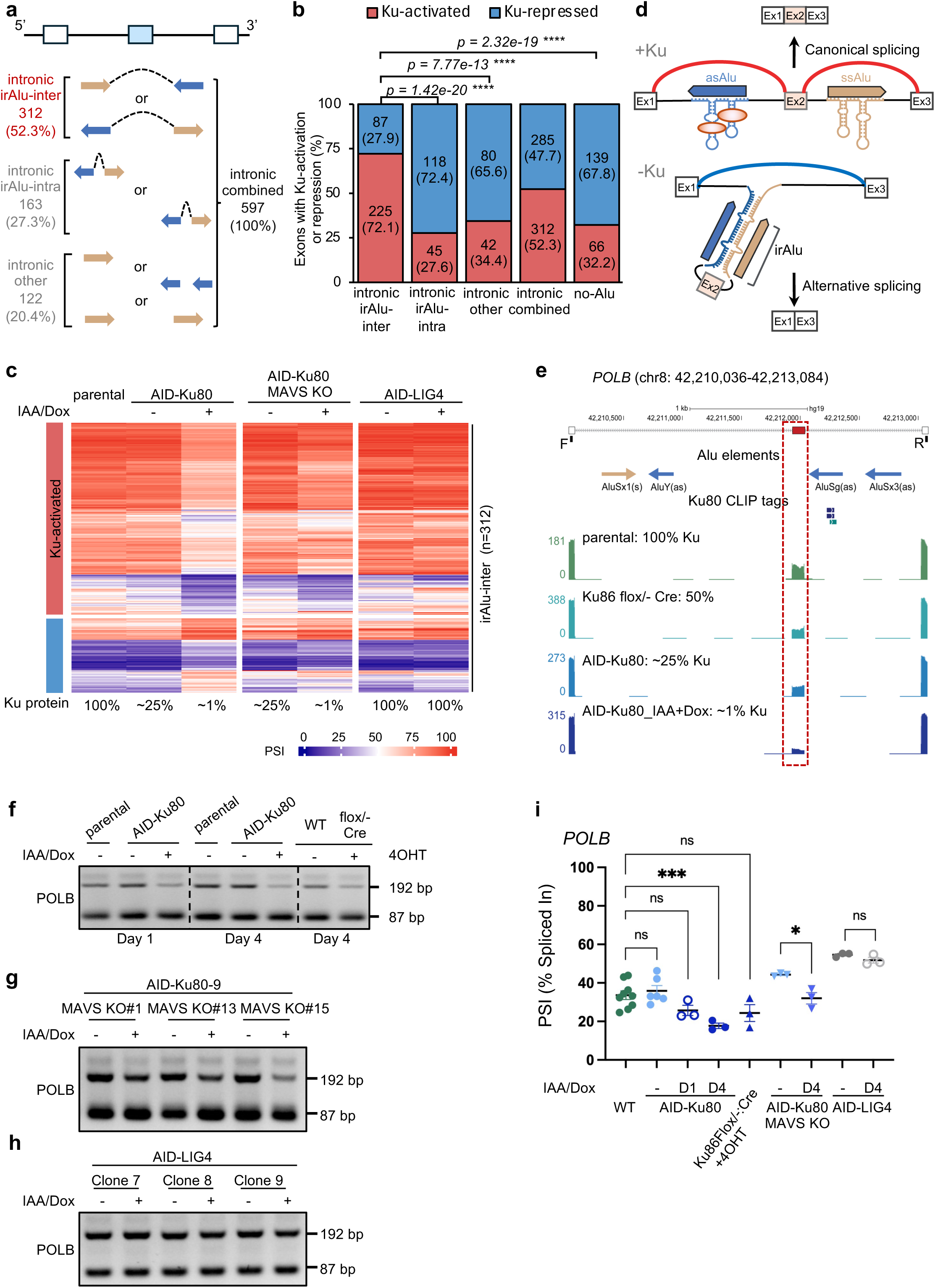
Ku directly represses skipping of exons flanked by intronic irAlu across the cassette exon. **a,** Schematics showing three mutually exclusive groups with different intronic Alu or Alu pair types. Cassette exons without exonic asAlu are classified as irAlu-inter if they contain at least one pair of opposite-orientated Alu elements located in the upstream and downstream intron, irAlu-intra if they contain at least one pair of opposite-orientated Alu elements located within the same intron (but not irAlu-inter), and other if they are neither two above groups: those containing only one intronic Alu element or same-oriented Alu elements located within one or both intronic regions. The relative abundance (percentage) of each group among high-confidence DSEs containing intronic Alu is also indicated. **b,** Percentage of Ku-repressed and Ku-activated high-confident DSEs grouped by type of intronic Alu elements contained: irAlu-inter, irAlu-intra, other, combined, no-Alu. Statistics showing bias towards Ku-activated or Ku-repressed exons associated with specific Alu-types are also indicated (Fisher’s exact test p<0.0001 ****; p>0.05 ns). **c,** PSI values for high-confidence Ku-repressed (blue) and Ku-activated cassette exons (red) containing intronic irAlu-inter in AID-Ku80 cells at 100%, ~25%, and ~1% Ku80 protein levels. PSI values in cells with MAVS KO; AID-Ku80 cells without or with Ku depletion (~25% and ~1% Ku) and in AID-LIG4 cells with or without LIG4 depletion are also shown. **d,** Schematic illustration of the proposed model by which Ku regulates cassette exons flanked by intronic irAlu-inter pairs. Ku binds monomeric asAlu, stabilizing stem-loop structures and inhibiting intronic irAlu pairing, thereby preventing exon looping out and promoting inclusion of the flanked exon. Upon Ku depletion, irAlu pairing induces exon looping and skipping. **e,** UCSC Genome Browser view of the *POLB* cassette exon (exon 6b) flanked by intronic irAlu-inter. Tracks show the positions of intronic sAlu and asAlu elements, Ku80-CLIP tags, and RNA-seq read coverage under varying Ku levels (100%, 50%, ~25%, and ~1%). **f-h**, Representative RT-PCR validation data for *POLB* transcripts in conditions described in Fig. 2 panels **(f-h)**. **i,** RT-PCR quantification of exon inclusion as percent-spliced-in (PSI) values from three independent biological replicates. Statistical significance of differential splicing was determined by the unpaired Student’s *t*-test *p*<0.0001 ****; *p*>0.05 ns.

Ku degradation induces exon skipping in 225 of 312 exons (72.1%) flanked by irAlu pairs (Fig. 3a-c), as compared to 32.2% of exons with noAlu (p=2.3e-19, Fisher’s exact test), or 27.6% exons flanked by irAlu-intra pairs in the upstream or downstream intron (p=1.4e-20, Fisher’s exact test; Fig. 3b). Among all cassette exons, irAlu-inter flanked exons are more likely to be activated by Ku, as compared to those without Alu (OR=2.3; Extended Data Fig. 3b). In any irAlu pair, at least one Alu element is in the antisense orientation. We previously showed that Ku binding sites preferentially resides at stem-loops within monomeric asAlu but are depleted at potential irAlu pairs, suggesting a model in which Ku inhibits irAlu pairing and thereby prevents irAlu-driven exon looping-out and skipping by stabilizing monomeric asAlu stem-loops (Fig. 3d). Consistent with this model, the 105-nt *POLB* exon 6b (35 aa) is included in Ku-proficient cells but skipped upon Ku loss (PSI changed from 29 to 13 as Ku falls from 100% to ~1%) (Fig. 3e). Robust Ku irCLIP signal was detected at the proximal downstream asAlu (AluSg), likely restricting its pairing with the upstream sAlu (AluSx1) (Fig. 3e). RT-PCR confirmed that Ku prevents *POLB* exon6b skipping in both AID-Ku80 and *Ku86^flox/-^:Cre-ERT2* cells (Fig. 3f,i). Although *POLB* is highly conserved (99% identity among primates; 87% human-rats), *POLB* alternative splicing variants are exceptionally prevalent in human cells (affecting ~half of all transcripts) but not seen in other mammalian and non-mammalian species^39^. Moreover, in three additional high-confidence Ku-dependent cassette exons flanked by irAlu, Ku reproducibly repressed exon skipping in both AID-Ku80 and *Ku86^flox/-^:Cre-ERT2* cells (Extended Data Fig. 4c).

As with asAlu-derived exons, Ku likely represses irAlu-induced exon skipping by directly binding the intronic asAlu within irAlu-inter pairs. This is supported by: (1) the presence of Ku irCLIP signal on one of the potentially involved asAlu elements (Fig. 3e and Extended Data Fig. 2d); (2) the correlation between PSI increase and Ku abundance (Fig. 3c,e,f,i); and (3) the Ku-dependent PSI changes are independent of G1 cell-cycle arrest (Extended Data Fig. 5a-c,f-g), dsRNA-/MAVS-mediated IFN signaling (Fig. 3g, i and Extended Data Fig. 6a,c) and the NHEJ pathway (evidenced by LIG4 degradation, Fig. 3h, i and Extended Data Fig. 6b,c). Yet, in contrast to the Ku-repressed and asAlu-derived “new” exons, the exons activated by Ku through suppressing irAlu-inter are in general deeply conserved across mammals, with a subset extending their conservation into vertebrates (Extended Data Fig. 7b). Thus, in addition to repressing asAlu-derived novel exons, Ku also safeguards pre-existing, evolutionarily conserved exons flanked by irAlu-inter pairs by preventing their aberrant primate-specific skipping.

Moreover, unlike asAlu-derived exons, 95% of which behaved as predicted by our model, 27.9% of irAlu-inter–flanked exons were instead repressed by Ku, in opposite to the model prediction. This might be explained by the presence of competitive irAlu pairs within the same intron. Prior work showed that inverted Alus within the same intron (irAlu-intra) can compete with intronic irAlu-iter pairing^6^. If the competing Alu is also in the anti-sense orientation, Ku binding to the asAlu in the competing irAlu-intra pairs would promote irAlu-inter formation, diluting the effect of Ku binding on the asAlu within the irAlu-inter (Extended Data Fig. 8a). To test this hypothesis, we subdivided Ku-dependent exons flanked by irAlu-inter based on the presence or absence of potential competitions (Extended Data Fig. 8a,b). Among the 57 irAlu-inter flanking cassette exons without competing irAlu-intra (intronic irAlu-inter only), 91% were activated by Ku (decreased PSI upon Ku depletion), a fraction significantly higher than those harboring both irAlu-inter and irAlu-intra (67.8% activated; p = 2.5e-4, Fisher’s exact test; Extended Data Fig. 8b), supporting a competing role of irAlu-intra in splicing regulation by preventing irAlu-inter formation.

Similar impact of competition can also be seen for irAlu-inter in the context of asAlu-derived exons (Extended Data Fig. 8c). Specifically, 97.1% (102/105) exonic asAlu without flanking irAlu-inter (exonic asAlu only) were repressed by Ku, as compared to 83.9% (26/31) among those containing irAlu-inter (exonic asAlu irAlu-inter flank; p=0.015, Fisher’s exact test; Extended Data Fig. 8d).

Overall, both the magnitude and consistency of Ku-dependent splicing changes are markedly greater for exonic asAlu in comparison to irAlu-inter. This might reflect the direct steric hinderance through direct binding of Ku on exonic asAlu. In contrast, the impact of Ku on exons flanked by irAlu-inter can be modulated by competing irAlu-intra, the distance between paired intronic irAlu elements, as well as other potential RNA- or Alu-binding proteins. Collectively, our findings suggest that about 40% of Ku-dependent exons can be explained by direct interaction of Ku with Alu elements via repressing asAlu-derived exons (149/967, 15.4%) or activating pre-existing exons flanked by irAlu (225/967, 23.2%) (Fig. 2b and 3b). In both cases, Ku suppressed the expression of evolutionarily new alternative splice variants specific for primates caused by Alu insertion, which are potentially deleterious.

### Ku has distinctive impact on cassette exon splicing from hnRNP C and DHX9

To understand how Ku’s impact on splicing compares to other known regulators of Alu-mediated alternative splicing and RBPs in general, we analyzed alternative exons regulated by other splicing factors, including hnRNP C, DHX9, PTBP1, and RBFOX2 using the same bioinformatics pipeline and statistical criteria^23,24,40^. Depletion of these splicing factors resulted in significant splicing changes (inclusion or exclusion) of 287-1,571 cassette exons (5.5-15.8% of all quantifiable cassette exons; Fig. 4a,b and Extended Data Table 3), which are less than or comparable to the number of exons affected by Ku depletion (~ 967), underscoring a broad impact of Ku in splicing regulation in human cells.

**Figure 4:**
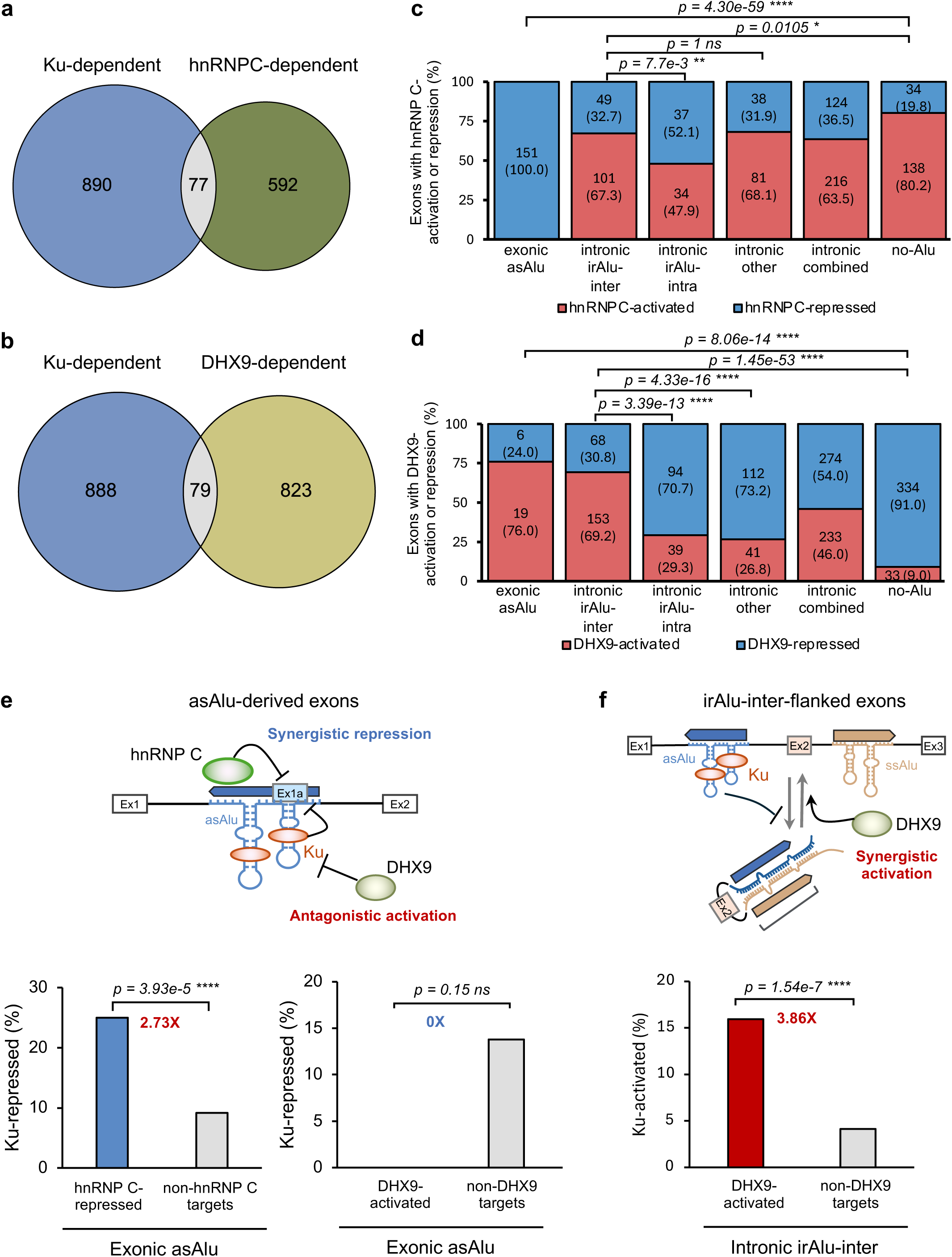
Distinct and combinational regulation of Ku-dependent splicing with hnRNP C and DHX9. **a,b,** Overlap between high-confidence Ku-dependent cassette exons with hnRNP C- (**a**) or DHX9-dependent exons (**b**). **c,d,** Percentage of hnRNP C-repressed and -activated cassette exons (**c**) as well as DHX9-repressed and-activated cassette exons (**d**) grouped by type of Alu elements contained: exonic asAlu, intronic irAlu-inter, intronic irAlu-intra, intronic other, intronic combined, and no-Alu. Statistics showing bias towards Ku-activated or Ku-repressed exons associated with specific Alu-types are also indicated (Fisher’s exact test p<0.0001 ****; p>0.05 ns). **e**, Synergistic or antagonistic regulation of Ku-dependent splicing of asAlu-derived exons by hnRNP C and DHX9. Top: Schematic illustration of the proposed mechanism. hnRNP C binding to polyU tracks upstream of asAlu-derived exons contributes to synergistic repression of exon inclusion with Ku, which stabilizes stem loops within asAlu. On the other hand, DHX9 destabilizes the stem loops through its RNA helicase function, which antagonizes Ku-dependent exon repression. Bottom: Percentage of Ku-repressed cassette exons among those derived from exonic asAlu with hnRNP C-dependent repression compared to those not regulated by hnRNP C (left) and percentage of Ku-repressed cassette exons among those derived from exonic asAlu with DHX9-dependent activation compared to those not regulated by DHX9. Odds ratios are indicated in color. **f**, Synergistic regulation of Ku-dependent splicing of irAlu-inter-flanked canonical exons by DHX9. Top: Schematic illustration of the proposed mechanism. DHX9 unwinds irAlu pairing through its RNA helicase function, which synergizes with Ku’s impact by stabilizing the stem loops of monomeric asAlu to inhibit irAlu pairing. Bottom: Percentage of Ku-activated exons among those flanked by intronic irAlu-inter with DHX9-dependent activation compared to those not regulated by DHX9. Statistical analysis: Fisher’s exact test p<0.0001 ****; p>0.05 ns (c-f).

Among these, hnRNP C and DHX9 are known regulators of Alu-mediated alternative splicing^23,24^. Among hnRNP C-dependent exons, exonic asAlu elements were exclusively suppressed by the presence of hnRNP C (151/151=100%), as compared to exons with no Alu (34/138=19.8%, p=4.3e-59, Fisher’s exact test; Fig. 4c), which recapitulates the previously reported results suggesting that hnRNP C competes with U2AF65 to block cryptic splice-site activation^23^ (Fig. 4c). In contrast, we found that among DHX9-dependent exons, both exonic asAlu and intronic irAlu-inter pairs were associated with exon inclusion (76% and 69.2%), as compared to 9% exons without Alu (p=8.06e-14 and p=1.45e-53, respectively, Fisher’s exact test; Fig. 4d). While Alu-mediated cassette exon skipping was not the focus of the original study^24^, our novel observation here is consistent with DHX9’s role as an RNA helicase that resolves long dsRNA structures to increase splice-site accessibility.

Moreover, Ku-regulated exons showed minimal overlap with the targets of hnRNP C or DHX9, as fewer than 8% of the 967 high-confidence Ku-dependent cassette exons intersected with either dataset (Fig. 4a,b), indicating that Ku acts through a mechanism distinct from hnRNP C and DHX9. Despite this independence, analysis of co-regulated exons revealed principles of combinatorial control. Specifically, asAlu-derived exons repressed by hnRNP C were 2.73-fold more likely to be also repressed by Ku, as compared to those not regulated by hnRNP C (p = 3.94e–5, Fisher’s exact test; Fig. 4e), indicating synergistic splicing repression. Similarly, irAlu-inter-flanked exons activated by DHX9 were 3.86-fold more likely to be activated by Ku than those irAlu-inter not regulated by DHX9 (p = 1.54e–7, Fisher, exact test; Fig. 4f), indicating synergistic splicing activation. Conversely, Ku and DHX9 appeared to exhibit a counteractive effect on the small subset of asAlu-derived exons activated by DHX9 (p=0.15, Fisher’s exact test; the moderate significance is due to the small number of asAlu-derived exons activated by DHX9; Fig. 4e), potentially by stabilizing (Ku) and unwinding (DHX9) the asAlu stem-loops. Together, these observations establish Ku as a broad acting and mechanistically distinct splicing regulator that works along the side with hnRNP C and DHX9 to safeguard splicing fidelity and efficiency, and especially in managing Alu-associated splicing challenges.

### Reduced Ku expression in the human brain correlates with higher expression of Alu-associated alternative splice variants

Next, we investigated the physiological consequences of Ku-regulated alternative splicing. While Ku is essential, we and others showed that 50% or even ~25% of Ku is compatible with human cell viability in tissue culture models (Fig. 1a-c and Extended Data Fig. 1a-d)^5,30,36^, allowing us to test whether Ku at ~50% or 25% levels can modulate splicing independent of viability. Using the same pipeline and stringent cut-off, we detected 891 and 902 cassette exons showing altered splicing between 100% vs. 50% Ku in the untreated *Ku86^flox/-^:Cre-ERT2* cells and between 100 vs. ~25% Ku in the untreated AID-Ku, respectively (Fig. 1c, Extended Data Fig. 9a,b). Among Ku-dependent exons in each case, 65-73% asAlu-derived exons were repressed by Ku whereas ~72% exons with flanking intronic irAlu-inter pairs were activated by Ku (p=3.84e-5 and 9.05e-12, respectively, Fisher’s exact test; Extended Data Fig. 9a,b), which is consistent with predictions of our models (Fig. 2c and 3c) and supports a dose-dependent role of Ku in splicing regulation on a significant subset of exons beyond cellular viability.

Alu-mediated tissues-specific alternative splicing has long been noted, and dysregulation of these events contributes to multiple neurological disorders^1^. In this context, we noticed that the mRNA levels of *XRCC5* (Ku80) and *XRCC6* (Ku70) are ~50% lower in bulk tissues derived from various brain regions compared to other non-brain tissues, based on GTEx RNA-seq data^41^ (Fig 5a and Extended Data Fig. 10). Together with dose-dependent splicing changes observed upon partial Ku depletion, this finding led us to propose a rate-limiting Ku hypothesis, which predicts that Ku-dependent exons may show differential splicing in the brain as compared to other tissues. To test this hypothesis, we identified a subset of 443 Ku-regulated exons showing differential splicing between brain vs non-brain tissues (FDR < 0.05, |ΔPSI|≥3) (Extended Data Table 6). Interestingly, among Ku-repressed exons derived from asAlu, 61.8% displayed higher inclusion in the brain, suggesting that these exons, normally repressed by Ku in other non-brain tissues, become preferentially included in brain tissue where Ku-dependent regulation might be rate-limiting (Fig. 5a,b). Conversely, for Ku-activated exons flanked by irAlu-inter, 68.9% demonstrate lower inclusion in the brain, again consistent with reduced Ku levels (Fig. 5a,b). The preference for higher or lower exon inclusion for these two groups of exons is statistically significant when compared to each other (p=0.0022, Fisher’s exact test), or to the “other” exons differentially spliced in the brain but not regulated by Ku, serving as a control, which showed a more balanced distribution between higher and lower inclusion in the brain (p=0.0227 and 0.0291, respectively, Fisher’s exact test; Fig. 5b). Two examples of such exons in *U2AF1L4* and *PALM*, and the PCR validation of tissue-specific splicing are shown in Fig. 5c,d. *U2AF1L4* (U2AF26) is a paralog of *U2AF1* (U2AF35) that can regulate both constitutive and alternative splicing^42^. Inclusion of the Ku-repressed exon overlapping with asAlu as the penultimate exon causes a frame shift that changes the 3’-terminal peptides of the encoded protein (Fig. 5e). *PALM* encodes a member of the paralemmin protein family implicated in plasma membrane dynamics in neurons and other cell types^43^. Here, Ku represses the skipping of a 132-nt exon (44 aa) flanked by irAlus, which has particularly high inclusion in the brain (Fig. 5f). Because Ku normally restrains “aberrant” Alu-driven splicing variants and human brain is known to have elevated dsRNA levels and hypersensitive to ADAR editing defects^44^, our data suggests that reduced Ku expression in the brain may provide a permissive environment that allow the emergence of otherwise silenced transcripts, often caused by Alu insertions (Fig. 5g).

**Figure 5:**
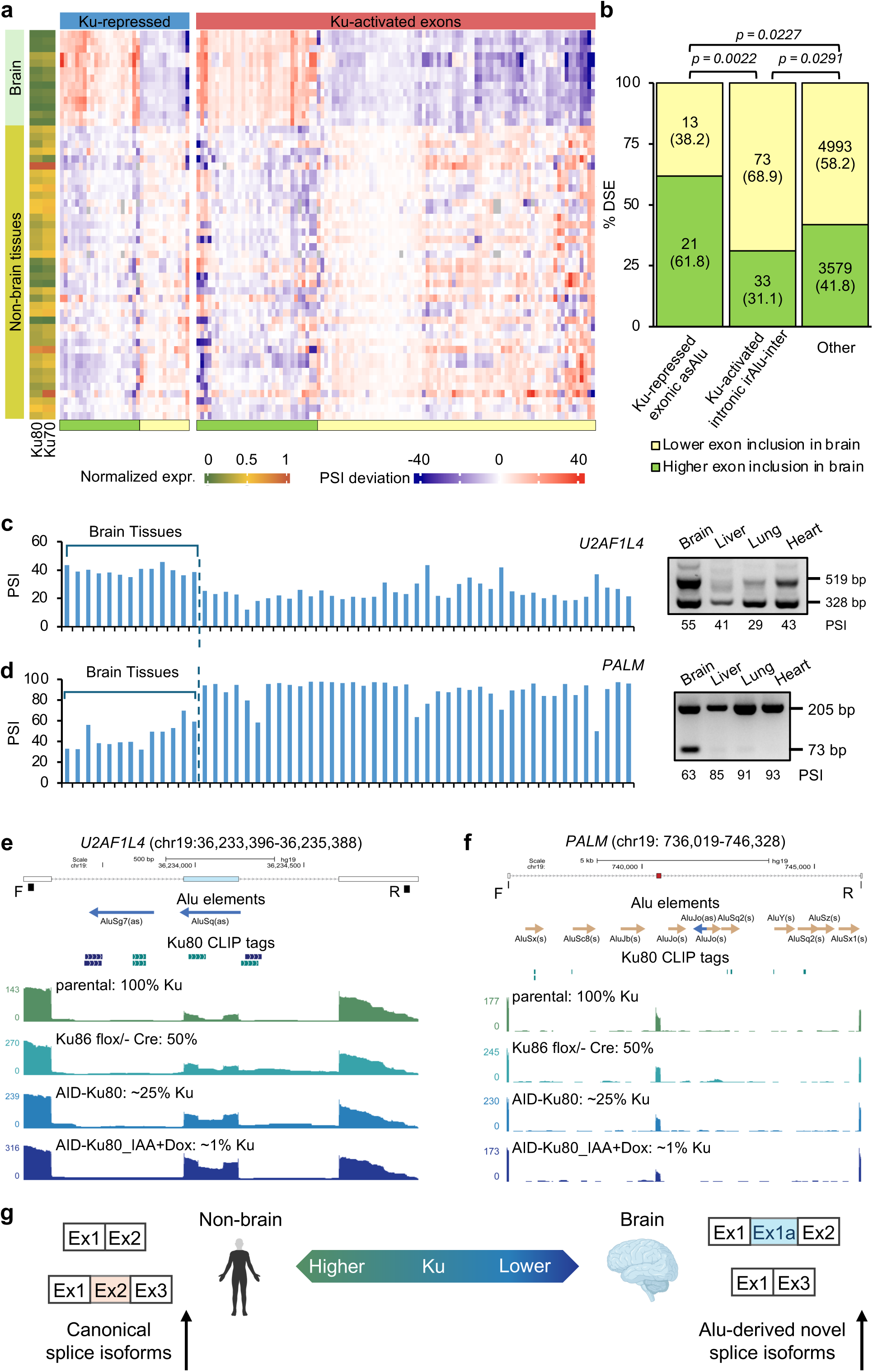
A lower Ku level in the brain is associated with higher expression of Alu-mediated alternative splice variants. **a,** Heatmap of normalized *XRCC5* (Ku80) and *XRCC6* (Ku70) mRNA expression (left) and cassette exon PSI deviation (middle and right) across various brain versus non-brain tissues quantified from GTEx RNA-seq data. Putative cassette exons under direct regulation by Ku (Ku-repressed exonic asAlu-containing and Ku-activated intronic irAlu-inter-containing high-confidence DSEs) and showing differential splicing between brain vs non-brain tissues were used for this analysis (**Method**). **b,** Proportion of high-confidence DSEs with higher or lower exon inclusion levels in brain vs. non-brain tissues among Ku-repressed exonic asAlu-containing and Ku-activated intronic irAlu-inter-containing DSEs. Other cassette exons differentially spliced between brain and non-brain tissues but not regulated by Ku are used for control. Statistics showing bias towards lower or higher exon inclusion in brain associated with specific types of cassette exons are also indicated (Fisher’s exact test p<0.0001 ****; p>0.05 ns). **c,d,** Two example cassette exons in *U2AF1L4* (**c**) and *PALM* (**d**) showing Ku dose-dependent regulation and differential splicing in the human brain. In each panel, PSI values across brain and non-brain tissues quantified from GTEx RNA-seq data is shown on the left. Representative RT-PCR gel images and corresponding PSI quantification from selected tissues are shown on the right. **e,f,** UCSC Genome Browser views for *U2AF1L4* (**e**) and *PALM* (**f**) showing the positions of Alu elements, Ku80-CLIP tags, and RNA-seq read coverages in cells with varying Ku levels (100%, 50%, ~25%, and ~1%). **g,** Schematic illustration of a proposed rate-limiting Ku-dependent splicing regulation model in which a lower Ku level in the brain provides a permissive environment that enables increased expression of novel splice variants introduced by Alu insertion.

### A heterozygous truncating mutation of Ku in a human patient is associated with developmental delay and neurological manifestations consistent with potential mitochondria dysfunction

Although pathogenic variants in most NHEJ factors (e.g., LIG4, DNA-PKcs) have been described in patients, causing radiosensitivity, severe combined immunodeficiency (SCID), and microcephaly, consistent with their role in DNA double strand break repair^45^, loss-of-function mutation in *XRCC5* or *XRCC6* is exceptionally intolerant in the human population (pLI = 1; LOEUF = 0.19 and 0.17, respectively). Given the dose-dependent splicing regulation by Ku, especially in brain-associated mRNA, and RNA-dependent innate immunity^5^—both relevant to neuronal function—and its naturally reduced expression in the brain, we reasoned that haploinsufficiency of Ku, if presents in patients, may cause neurological dysfunction. Through GeneMatcher^46^, we identified a 15-year-old girl carrying a *de novo* heterozygous *XRCC5*/Ku80 nonsense variant (c.421C>T; p.R141X). Introducing this mutation into human cells reduced *XRCC5*/Ku80 expression by ~50% (Extended Data Fig. 11a, b). Clinically, the patient exhibited early-onset developmental delay (3 months), microcephaly, speech delay, hypotonia, scoliosis, recurrent seizures, repeated viral pneumonias, and episodes of metabolic acidosis. Given that Ku regulates ~10% of quantifiable cassette exons, broad multisystem involvement is plausible, as observed from haploinsufficiency of other splicing regulators (e.g., hnRNP C^47^). We note that the lactic acidosis and neuromuscular defects are consistent with a prominent role of Ku in ensuing proper splicing of mitochondrial genes (Fig. 1e). Among the 967 high-confidence exons, 133 exons fall into 99 nuclear-encoded mitochondrial genes (Extended Data Table 5). Among them, 28 genes showed significantly reduced expression upon Ku depletion, including three IFN-independent targets—*MMUT*, *MTO1*, and *ATP5F1*. Both *MMUT* and *MTO1* harbor asAlu-derived exons, whereas ATP5F1 has a Ku-dependent exon flanked by irAlu. In *MTO1*, Ku degradation causes aberrant inclusion asAlu-derived exons (PSI from ~15% to >75%) (Extended Data Fig. 11c-d) at the cost of reduction in the canonical form and decrease of overall *MTO1* mRNA by ~50% (Extended Data Fig. 11e). MTO1 catalyzes the 5-carboxymethylaminomethyl modification of wobble uridine (U34) in multiple mitochondrial tRNA essential for mitochondrial translation^48^. Pathogenic variants in MTO1 cause lactic acidosis, developmental and neurologic abnormalities, overlapping with the XRCC5-deficiency^49,50^. These findings suggest that Ku haploinsufficiency may disrupt splicing in the brain, including in mitochondrial genes, offering a possible explanation for the neurological and metabolic (e.g., prone to acidosis) features observed in the patient.

## Discussion

This study establishes Ku as a major splicing regulator in humans and likely also other higher primates, through a mechanism distinct from previously characterized splicing factors. Whereas most splicing factors recognize short single-stranded RNA motif sites^8^, Ku forms a constitutive ring encircling dsDNA ends and dsRNA stem-loop structures in a sequence-independent and potentially structure-selective manner. A major class of Ku-bound RNAs are primate-specific Alu elements, which frequently reside in gene-dense regions and profoundly reshape transcriptomes evolution by introducing new splicing variants. The most notable manifestation of this process is the “exonization” of Alu elements^3,16,17^, which accounts for ~7% of annotated cassette exons in human genome, including ~6% derived from asAlu. Because many of these insertions disrupt the reading frame and they are frequently predicted to be deleterious, safeguarding mechanisms, such as hnRNP C^23^, play a critical role in preventing aberrant asAlu-derived exon inclusion. More broadly, the RNA splicing mechanism is optimized to distinguish *bona fide* exons from other intronic sequences and thereby avoiding random sequences from being recognized as exons^51^. Building on this framework, our work identified Ku as a novel regulator of alternative splicing, providing insights on the impact of Alu on primate evolution and the physiological function of Ku-RNA interaction in human diseases.

Here we show that Ku ring suppresses multiple modes of Alu-mediated alternative splicing. Beyond directly repressing 13.4% (149 of 1,112 quantifiable exons) asAlu-derived exons, Ku also prevents irAlu-mediated exon looping-out of 225 canonical exons flanked by irAlu, the largest class of Alu-mediated alternative splicing events in our study. While the contribution of Alu exonization to primate-specific splicing has long been recognized, irAlu-mediated skipping of canonical exons in primates has been more difficult to characterize in part owing to technical challenges in defining irAlu pairs and their numerous competing configurations^21^, leaving it unclear whether and how such events are regulated. In this context, our study uncovered a hidden layer of primate-specific transcriptomic regulation. In both cases, Ku suppresses the evolutionarily new isoforms, which might be deleterious, presumably through stabilizing stem loops within asAlu. We note that these 374 exons almost certainly represent an underestimate of Alu-mediated splicing regulated by Ku. This is because many Alu-mediated alternative splice variants have low abundance due to inherently weak splicing signals, so that the magnitude of Ku-dependent splicing might be minute and fall below stringent statistical cutoff used in this study (|ΔPSI| ≥ 10). It is important to emphasize that in contrast to other splicing factors previously known to regulate Alu-mediated splicing events, such as hnRNP C and DHX9, which themselves are highly conserved in mammals, Ku expression increased ~100-fold during primate evolution in parallel with Alu expansion, positioning it as a co-opted suppressor of evolutionarily new, frequently NMD-coupled splice variants generated by Alu insertion.

On the other hand, approximately 60% of Ku-dependent exons cannot be explained by exonic asAlu or flanking intronic irAlu pairs, which suggests additional mechanisms contributing to the remaining events, including the presence of competing Alu elements, Ku binding to non-Alu stem loops, as well as secondary effects such as regulating splicing of other splicing factor-encoding genes. Moreover, although we clearly demonstrate that Ku has a direct role in pre-mRNA splicing that is independent of the cell cycle and interferon/innate immune signaling, it remains an open question to what extent Ku-mediated splicing defects contribute to dsRNA-dependent innate immune activation. In particular, altered splicing may promote the accumulation of highly structured RNAs (e.g., asAlu-containing introns and pre-mRNA), thereby amplifying innate immune responses and reinforcing growth arrest.

Our findings also highlight the tissue-specific, functionally selective, and disease-relevant consequences of Ku-dependent regulation of Alu-mediated splicing. Although most of these events generate deleterious “leaky” products, a subset might have acquired adaptive value—for example, the irAlu-mediated skipping of *TBXT* exon 4 that contributed to tail loss in apes^22^. In this context, while *XRCC5* and *XRCC6* are among the most loss-intolerant human genes (pLI = 1), we show that reduced Ku level in the human brain correlates with increased expression of Alu-mediated alternative splice variants, and that Ku loss significantly affects the splicing of multiple mitochondrial genes, raising the intriguing possibility that such Ku-limited splicing programs could influence primate cognitive evolution or susceptibility to neurologic and neurodegenerative diseases. In line with this notion, we identified a patient with a *de novo* heterozygous loss-of-function variant in *XRCC5*/Ku80 who manifests severe neurodevelopmental deficits and metabolic acidosis, features consistent with preferential brain involvement and mitochondrial dysfunction. Notably, altered splicing of mitochondrial genes is increasingly linked to neurological diseases^1^. Finally, Ku binds not only Alu-derived RNAs but also a broad spectrum of other structured RNAs, including rRNAs, tRNAs, and microRNAs^5,31^, indicating that Ku might have acquired expanded RNA-regulatory functions during primate evolution. Together, these observations reveal Ku as both a genome guardian and a transcriptome safeguard, integrating Alu expansion, RNA processing, and primate-specific biology.

## Acknowledgement

We thank members of the Zha and Zhang laboratories at Columbia University for scientific input and helpful discussions during this study. We thank the patient and her family.

## Author contributions

S.Z. and C.Z. conceived the study; T.Y., J.Y., S.Z. and C.Z. designed the study; T.Y. performed most of the bioinformatics analysis with initial contribution from D.F.M.; J.Y. performed experimental validation; Y.Z. A.L. and B.J.L generated and characterized the AID-Ku80 and control AID-LIG4, and MAVS KO AID-Ku80 cells, collected the RNA-seq and performed differential expression analyses. Q.D, F.H., M.Y., V.A.G, K.Z, and L.C. contributed to the human genetic studies: F.H. and M.Y. generated HEK293T cell line carrying *XRCC5* c.421 C>T heterozygous mutations; Q.D performed sequencing and RT-PCR; D.J. and C.L. are the neurologists who identified this patient and her *XRCC5* mutation; V.A.G. helped with GeneMatcher. T.Y., J.Y., S.Z. and C.Z. wrote the manuscript. All authors critically reviewed and approved the final manuscript.

## Methods

### Cell culture

The HCT116 tet-OsTIR1 cells were purchased from RIKEN Cell Bank, cultured in McCoy’s 5A medium (Corning 10-050-CV) supplemented with 10% fetal bovine serum (Hyclone SH30071.03), 2 mM L-glutamine (Gibco 25030-081) and penicillin/streptomycin (Gibco 15140-122), and incubated at 37°C. Cell lines with AID tags added to both endogenous Ku80 allele via CRISPR-aided targeting were established previously^5^. To degrade Ku80, 0.5 μg/mL doxycycline (Dox, Sigma D9891) and 100 μM indole-3-acetic acid (IAA, Sigma I3750-5G-A) were added to the medium the day after seeding. Similarly, HCT116 *Ku86^flox/-^* and *Ku86^flox/-^;CreERT2* cells were previously characterized^5^. To deplete Ku in these cells, 4OHT at 50nM final concentration was used to induce Cre-nuclear translocation and Ku80 (also referred to as Ku86 in human cells) excision.

### RNA extraction and RT-PCR

Total RNA was extracted using TRIzol reagent (Thermo Fisher Scientific 15596018) according to the manufacturer’s instructions with minor modifications. Cells were lysed directly in 1 mL of ice-cold TRIzol and incubated for 5 min at room temperature. Phase separation was achieved by adding 0.2 mL chloroform, mixing vigorously, and centrifuging at 12,000 g for 15 min at 4 °C. The aqueous phase was transferred to a new tube, and RNA was precipitated with 0.5 mL cold isopropanol followed by incubation at 4 °C for 10 min and centrifugation at 12,000 g for 10 min at 4 °C. Pellets were washed twice with 75% ethanol, air-dried, and resuspended in nuclease-free water. RNA was solubilized for 10 min at room temperature and further incubated at room temperature with gentle agitation. RNA concentration and purity were measured using a NanoDrop spectrophotometer, and samples with A260/A280 ratios of ~2.0 were used for downstream applications.

For reverse transcription, total RNA was first treated with DNase I to remove residual genomic DNA. Up to 2 µg of RNA was incubated with DNase I in the presence of RNase inhibitor in a 10 µL reaction, followed by incubation at 25 °C for 30 min and heat inactivation at 75 °C for 5 min. DNase-treated RNA was then mixed with oligo(dT) primers and dNTPs, heated to 65 °C for 5 min to facilitate primer annealing, and rapidly cooled on ice. All cDNA were generated using SuperScript™ IV First-Strand Synthesis System (Invitrogen 18091050) in the supplied reaction buffer with DTT and RNase inhibitor, according to the manufacturer’s guidelines. Reactions were incubated at 50 °C for 30 min, followed by enzyme inactivation at 80 °C for 10 min. RNA templates were subsequently removed by adding RNase H and incubating at 37 °C for 20 min. The resulting cDNA was stored at –20 °C until use.

For PCR amplification, Taq DNA polymerase (Qiagen, P7050L) was used following the manufacturer’s instructions. Briefly, cDNA templates were amplified in standard Taq buffer containing MgClL, dNTPs, and cassette exon-specific primers. Reactions were denatured at 94 °C, subjected to 35 cycles of amplification (94 °C for denaturation, primer-specific annealing temperature for 30 s, and 30 – 45 s long incubation at 72 °C for extension), and completed with a final extension at 72 °C for 5 min. PCR products were analyzed on 2% agarose gels stained with ethidium bromide. All DNA oligos used in this study are summarized in Extended Data Table 7.

### RNA-seq data generation and analysis

For transcriptome-wide profiling of Ku-dependent splicing, we used the AID system including parental controls treated with DMSO or IAA/Dox at two time-points (day 2 and 4), serving as 100% Ku protein controls. Ku80-AID-tagged cell lines treated with DMSO showed partially reduced Ku expression (~25% Ku), while co-treatment of IAA/Dox for four days resulted in near-complete depletion (~1% Ku protein). This design enabled precise titration of intracellular Ku levels and allowed for high-resolution mapping of differential splicing events across different Ku levels. As a second system with genetic depletion of Ku for comparison, we also profiled splicing in *Ku86 ^flox/-^* or *Ku86 ^flox/-^;CreERT2* HCT116 cells treated with Dox to represent 50% and 0% Ku expression, respectively. All conditions were sampled in biological triplicates (except for *Ku86^flox/-^; CreERT2* cells, which have two replicates). Standard paired-end RNA-seq libraries for control and Ku depleted samples were prepared using poly-dT (for all Day4 treatment used in initial analyses sequenced by GeneWiz). To determine the immediate impact of Ku-deletion (24hr) or the contribution of cell cycle (CDK4/6 inhibitor treated for 48hr), AID-Ku clones were induced with IAA/Dox for 24hr or together with CDK4/6 inhibitor for 48hr as detailed previously. The total RNAs were collected and sequenced using RiboZero protocols and on Ilumina NovaSeq 6000 at Columbia Genome Resource Center. For comparison, we also downloaded published RNA-seq data upon depletion of hnRNP C^23^, DHX9^24^, PTBP1^40^, RBFOX2^40^ and their respective controls. A summary of RNA-seq datasets analyzed in this study is provided in Extended Data Table 1.

We processed and analyzed RNA-seq data using the Quantas pipeline, as described previously (https://zhanglab.c2b2.columbia.edu/index.php/Quantas_Documentation)^34^. Briefly, we mapped paired-end RNA-seq reads to the reference transcriptome and exon junctions (GRCh47/hg19) with OLego^35^. We then used mapped reads to infer the transcript structures, followed by quantifying gene expression in Reads Per Kilobase of transcript per Million mapped reads (RPKM) and exon inclusion values in percent spliced in (PSI). Statistical analysis was performed to identify differentially spliced events (DSEs) using Fisher’s exact test across six major classes of AS events (cass, taca, alt3, alt5, iret, and mutx) between each pair of conditions (Extended Data Table 2). Exons satisfy the following criteria are considered as DSEs: 1) read coverage ≥ 20, 2) |ΔPSI| ≥ 10, and 3) FDR ≤0.05. An exon with junction read coverage ≥20 is called quantifiable and only quantifiable exons were used for differential splicing analysis. Differential expression was performed using DESeq2 (ref.^52^).

For cassette exons, high-confidence Ku-dependent DSEs were defined as those showing consistent splicing changes in both 100% vs. ~1% Ku and ~25% vs. ~1% Ku comparisons, which were used for detailed analysis of splicing-regulation mechanisms. For hnRNP C and DHX9, high-confidence DSEs were defined as those showing consistent direction of changes upon RBP depletion using two independent shRNAs, similar to the original studies^23,24^. For PTBP1 and RBFOX2, high-confidence DSEs were defined as those showing consistent direction of changes in K562 and HepG2 cells (Extended Data Table 3).

### Annotation of Alu elements

We downloaded the genomic coordinates of repeat masked regions from the UCSC Genome Browser (i.e., the RepeatMasker track)^53^, from which Alu elements were extracted. Custom python scripts were then used to compare coordinates of cassette exons and Alu elements to classify exonic and intronic Alu in the sense or antisense strand. We required exonic asAlu and sAlu to have ≥10 bp overlap with the cassette exons, whereas intronic asAlu or sAlu elements are completely contained within flanking upstream or downstream introns. To determine irAlu pairs, we only considered the orientation of the intronic Alu elements and did not impose any distance threshold, which does not appear to improve signal-to-noise ratio. Different types of irAlu pairs include those spanning the cassette exon (irAlu-inter) and those within introns (irAlu-intra).

### Exon conservation across primate and other species

Multiple genomic sequence alignments of 100 species, including human, were downloaded from the UCSC genome browser^53^. For each cassette exon, we extended 200 nt upstream and 200 nt downstream and then extracted its aligned orthologous sequences. To determine if this exon is conserved in each of the other 99 species, we checked whether the splice site dinucleotides GT/AG are conserved. The exon was called conserved if both 3’ and 5’ splice sites of the exon are conserved.

### Ku expression and differential splicing across human tissues

We downloaded gene transcripts per million (TPM) values, exon junction read counts (GTEx Analysis V8) of 17,382 RNA-sequencing samples from 838 human adults at the GTEx Portal (https://gtexportal.org/) (The GTEx Consortium 2020). Tissue-level gene expression values were calculated by averaging all samples within each tissue type (13 brain and 40 non-brain tissues). *XRCC5* (Ku80) and *XRCC6* (Ku70) tissue-level gene expression levels were extracted and then min-max normalized to the range of 0 to 1 for cross-tissue comparisons through the following formula: normalized tissue-level TPM = (*x*-*min*)/(*max*-*min*), where *x* is tissue-level TPM, *min* and *max* are the minimum and maximum tissue-level TPM for *XRCC5* or *XRCC6*.

Cassette exon PSI values across human tissues were quantified using the Quantas pipeline^34^. Briefly, we first converted the coordinates of exon junctions from GTEx from hg38 to hg19 using liftOver^54^. These exon junction coordinates were then matched against the exon junctions of the inclusion or skipping isoforms of each cassette exon and the PSI value was calculated using the matched exon-junction read counts. Only cassette exons with both inclusion junctions and the skipping junction matched were included in this analysis. Read counts from individuals within each tissue type were pooled together for the estimation per tissue. The mean PSI across all tissues was then subtracted to calculate PSI deviation for each tissue. To identify cassette exons with differential splicing patterns between brain and non-brain tissues, we performed a two-sided t-test between brain and non-brain tissue-level PSI values. Cassette exons with Benjamini-Hochberg corrected FDR ≤ 0.05 and |ΔPSI| ≥ 3 between brain and non-brain tissues, and ≤5 tissues missing PSI vales are defined as brain-specific (Extended Data Table 6).

### Gene Ontology analysis

We performed gene ontology (GO) term enrichment analysis with DAVID^55^ for genes containing high-confidence Ku-dependent DSEs. GO terms with a Benjamini FDR ≤0.05 were reported in this study. The lists of genes associated with “RNA-binding”, “DNA repair” and “mitochondrion” are downloaded and provided in Extended Data Table 5.

### Whole exome sequencing of *XRCC5*-deficient patient

DNA derived from somatic tissue was subjected to whole-exome sequencing (WES). Trio-based WES was performed by Running-gene (Beijing, China), including exome library preparation, sequencing, and data analysis. Libraries were generated using SureSelect Human All Exon V6. The Illumina NovaSeq 6000 (Illumina, SanDiego, CA, United States) with PE 150 was used to perform subsequent sequencing, with depth over 100x. Reads were aligned to the hg19 version of the human genome.

### Generating HEK293T knockin cells carrying *XRCC5* c.421 C>T (p. R141X) mutation

The HEK293T cell line with the patient XRCC5 c.421 C>T (p. R141X) mutation was generated using a previously described method^56^ with modifications. Briefly, HEK293T cells were transfected with the pLentiCRISPR V2-XRCC5-guide RNA (sgRNA) plasmids and the donor vector with the patient mutation. In a parallel set of experiments to generate the control HEK293T cell line, the pLentiCRISPR V2 empty vector was used in transfection. Two days after transfection, cells were treated with puromycin (2 μg/μl; Sigma-Aldrich) and maintained in the same media for up to 2 weeks. Single cells were plated, and individual single–cell–originated clones were propagated and examined to select clones carrying the targeted *XRCC5* mutation.

The following sgRNAs were used: sgRNA#1 (5′-TCTCCAGAGATACACTTCCA-3′) and sgRNA#2 (5′-AGGATCTTACAAGAATTGCA-3′).

## Data availability

All data generated or analyzed during this study are included in this article (and its supplementary information and Source Data files). Ku RNA-seq data will be deposited to NCBI/GEO. Published RNA-seq data used in this study were downloaded from different repositories: hnRNP C in HeLa cells (ArrayExpress: E-MTAB-1147) and DHX9 in HEK293 cells (GEO: GSE85164), RBFOX2 and PTBP1 in HepG2 and K562 cells (ENCODE Data Coordination Center: ENCSR456FVU) (also see summary in Extended Data Table 1).

## Extended Data Figure Legends

**Extended Data Figure 1: Ku depletion results in global changes in different types of alternative splicing.**

**a,** Western blot analysis validating efficient depletion of AID-tagged Ku80 in HCT116 cells at day 1 (D1) and day 4 (D4) of IAA and Dox treatment. **b,** Distribution of DSEs detected in 100% vs. ~25% Ku, 100% vs. ~1% Ku, and ~25% vs. ~1% Ku, degraded in AID system, with respective to six major classes of alternative splicing events: alternative 3’ or 5’ splice-site selection, cassette exons, intron retention, mutually exclusive exon, and tandem cassette exons. **c,** Western blot analysis showing Ku depletion in *Ku86^flox/-^;CreERT2* HCT116 cells upon 4OHT treatment. **d,** Distribution of DSEs detected in 50% vs. ~1% Ku, degraded in Cre/Flox system, with respective to the same six major classes of alternative splicing events as in (b). **e,** Overlap of cassette exons (CEs) activated or repressed by Ku between 100% vs. ~1% Ku and ~25% vs. ~1% Ku comparisons. **f,** Comparison of ΔPSI (change in Percent Spliced In) between 100% vs. ~25% Ku and 100% vs. ~1% Ku (left) as well as ~25% vs. ~1% Ku and 100% vs. ~1% Ku (right). In each panel, cassette exons with differential splicing in either comparison are plotted. Number and percentage of DSEs common to both conditions are indicated in labels.

**Extended Data Figure 2: Example Ku-dependent cassette exons involved in RNA-binding, mitochondrial function, and DNA repair.**

**a,** UCSC Genome Browser view of the *COQ3* cassette exon containing asAlu with no flanking irAlu elements. Tracks show the positions of intronic sAlu and asAlu elements, Ku80-CLIP tags, RNA-seq read coverage under varying Ku levels (100%, 50%, ~25%, and ~1%), and conservation. **b,** Similar to (**a**) but UCSC Genome Browser view of the *MTO1* cassette exon containing asAlu with no flanking irAlu pars. **c,** Similar to (**a**) but UCSC Genome Browser view of the *MBNL2* cassette exon containing intronic irAlu-iter with competing intronic irAlu-intra elements. **d**, Similar to (**a**) but UCSC Genome Browser view of the *RAD1* cassette exon containing intronic irAlu-iter elements. Note FLAM_A, or free left Alu monomer (FLAM) in the upstream intron is an ancient form of Alu, which is likely involved in irAlu pairing given the absence of other Alu elements in the upstream intron.

**Extended Data Figure 3: Ku-dependent DSEs are more enriched among cassette exons containing exonic asAlu or those flanked by intronic-irAlu-inter pairs.**

Percentage of Ku-activated and Ku-repressed cassette exons among all quantifiable cassette exons containing different types of Alu elements: exonic asAlu or sAlu, intronic Alu, and no Alu (a); intronic irAlu-inter or irAlu-intra, other intronic Alus, intronic Alu combined, and no Alu (b).

**Extended Data Figure 4: RT-PCR validation of Ku-dependent cassette exons.**

**a,** Representative gel images of RT-PCR validation for *NUP50*, *TTLL5*, and *RPE* transcripts containing exonic asAlu elements. **b,** The sequence of the *EXOSC9* alternative exon-derived from asAlu. The cryptic splicing sites and the exon (underlined) are marked together with the corresponding amino acid sequence. **c.** Representative gel images of RT-PCR validation data for *NUP88*, *APAF1*, and *DONSON* transcripts flanked by intronic irAlu elements. More details of analyzed conditions are described in Fig. 2 panels (f-i) in the main text.

**Extended Data Figure 5: Ku directly represses Alu-mediated alternative splicing upon Ku depletion at day 1 or in G1 arrest cells.**

**a,** Distribution of 379 Ku-dependent cassette exons detected in cycling HCT116 cells upon Ku depletion at day 1 or G1-arrest HCT116 cells at day 2, with respective to type of Alu elements contained. DSEs showing statistically significant differential splicing in at least one comparison and consistent directions of splicing changes between the two comparisons are included in the analysis. **b,** Percentage of Ku-repressed and Ku-activated DSEs grouped by type of Alu elements contained: exonic asAlu, intronic irAlu-inter, intronic irAlu-intra, intronic other, intronic combined, and no-Alu. For each Alu-type, the number and percentage (in parenthesis) of exons are indicated. Statistics showing bias towards lower or higher exon inclusion in brain associated with specific types of cassette exons are also indicated (Fisher’s exact test p<0.0001 ****; p>0.05 ns). **c,** Heatmap showing PSI values for Ku-activated intronic irAlu-inter (red) and Ku-repressed exonic asAlu-containing (blue) DSEs in cycling AID-Ku80 HCT116 cells without or with Ku depletion at day 1 as well as G1 arrested AID-Ku80 HCT116 cells without or with Ku depletion at day 2. Within cassette exons for analysis, those with FDR ≤ 0.05, PSI ≥ 10 or PSI ≤ −10 in either condition are defined as Ku-activated and -repressed DSEs. **d,** UCSC Genome Browser view of the *EXOSC9* cassette exon containing an exonic asAlu element. Tracks show the positions of intronic sAlu and asAlu elements, Ku80-CLIP tags, RNA-seq read coverages in both cycling cells at day 1 and G1 arrest cells at day 2, with or without IAA and Dox treatment to induce Ku depletion. **e,** Representative gel images showing RT-PCR validation for *EXOSC9* transcripts in parental and AID-Ku80-expressing HCT116 cells treated with CDK4/6 inhibitors for 48 h, with or without IAA and Dox treatment for Ku depletion, with PSI quantification shown at the bottom. **f,** UCSC Genome Browser view of the *POLB* cassette exon containing an exonic asAlu element. Tracks show the same information as (d). **g,** Representative gel images showing RT-PCR validation for *POLB* transcripts in parental and AID-Ku80-expressing HCT116 cells treated with CDK4/6 inhibitors for 48 h, with or without IAA and Dox treatment for Ku depletion, with PSI quantification shown at the bottom. For all RT-PCR quantification of exon inclusion as PSI values, three independent biological replicates were used. Statistical significance of differential splicing was determined by the unpaired student’s *t*-test *p*<0.0001 ****; *p*>0.05 ns.

**Extended Data Figure 6: Ku regulates alternative splicing independent of NHEJ requiring Lig4 and interferon (IFN) and NF-κB signaling requiring MAVS.**

**a,** Western blot analysis of pSTAT1, STAT1, MDA5, RIG-I, MAVS and Ku80 upon IAA/Dox-induced Ku depletion on day 4. **b,** Western blot analysis confirming efficient depletion of AID-tagged Lig4 in HCT116 cells at day 4 after IAA and Dox treatment. Quantification of all protein levels normalized to Vinculin from two independent western blot replicates. Y-axis is displayed on a log scale for Lig4 and Ku80. **c,** PSI values for 967 high-confident DSEs in AID-Ku80 cells at 100%, ~25%, and ~1% Ku80 protein levels. PSI values in MAVS KO; AID-Ku80 cells without or with Ku depletion (~25% and ~1% Ku) and AID-LIG4 cells with or without LIG4 depletion are also shown.

DSEs were grouped by type of Alu elements containing: exonic asAlu, intronic irAlu-inter, irAlu-intra, other, no Alu, and exonic sAlu.

**Extended Data Figure 7: Conservation of Ku-regulated exons.**

**a,** Ku-repressed exons derived from asAlu. **b,** Ku-activated exons flanked by intronic irAlu-inter. In each panel, the heatmap shows conservation of exons in each of the 100 vertebrate species, with conserved exons indicated in green. Data from human are shown in the first column.

**Extended Data Figure 8: Competition of different types of Alu elements in Ku-dependent splicing.**

**a,** Schematics showing the classification of two intronic irAlu-inter groups: intronic irAlu-inter only if no irAlu-intra presents or intronic irAlu-inter&intra if irAlu-intra presents. The relative abundance (percentage) of each group among high-confidence Ku-dependent cassette exons is also indicated. **b,** Percentage of Ku-repressed and Ku-activated high-confidence DSEs grouped by intronic irAlu type: intronic irAlu-inter only, intronic irAlu-inter&intra, and intronic irAlu-intra, other, combined, and no Alu controls. The number and percentage (in parenthesis) of exons in each group are also indicated. Statistics showing bias towards Ku-activated or Ku-repressed exons associated with specific Alu-types are also indicated (Fisher’s Exact Test *p*<0.0001 ****; *p*>0.05 ns). **c**, Schematics showing the classification of three exonic asAlu groups: exonic asAlu only if they are not flanked by intronic irAlu-inter, exonic asAlu (irAlu-inter flank) if they are flanked by intronic irAlu-inter only, or exonic asAlu (irAlu-inter&intra flank) if they are flanked by both intronic irAlu-inter and irAlu-intra. **d,** Percentage of Ku-repressed and Ku-activated high-confidence DSEs grouped by exonic asAlu type: exonic asAlu only, exonic asAlu (irAlu-inter&intra flank), exonic asAlu (irAlu-inter flank), and no Alu control. The number and percentage (in parenthesis) of exons in each group are also indicated. Statistics showing bias towards Ku-activated or Ku-repressed exons associated with specific Alu-types are also indicated (Fisher’s Exact Test).

**Extended Data Figure 9: A subset of Ku-dependent cassette exons shows Ku dose-dependent splicing changes.**

**a,** DSEs between 100% vs. ~25% Ku. **b,** DSEs between 100% vs. 50% Ku. In each panel, percentage of Ku-repressed and Ku-activated DSEs grouped by Alu types, exonic asAlu and intronic irAlu-inter, are shown. The breakdown of all DSEs is also shown for comparison. The number and percentage (in parenthesis) of exons are indicated for each bar.

**Extended Data Figure 10: Violin plots showing the distribution of *XRCC5* (Ku80) and *XRCC6* (Ku70) mRNA levels (TPM) across human tissues.** These Violin plots were generated using GTEx Data Portal.

**Extended Data Figure 11: *XRCC5* heterozygous loss-of-function mutation in a human patient with neurological manifestations and mitochondrial function deficiency. a,** Scheme showing the truncating mutation derived from patients and targeted into HEK293T cells. **b,** qPCR shows the mRNA levels reduced to 50% in heterozygous targeted cells, as compared to control. c-e, Ku depletion leads to aberrant splicing and expression of MTO1, which modifies multiple mitochondrial tRNA essential for mitochondrial translation. **c**, Genome Browser view of the *MTO1* cassette exons derived from asAlu This exon is skipped in the canonical form but included as two variants due to alternative 3’ splice sites with or without retention of the upstream intron, generating multiple alternative splice isoforms. **d,** RT-PCR validation of the *MTO1* asAlu-derived exon splicing with or without intron retention upon Ku degradation. More details of analyzed conditions are described in Fig. 2 panels (f-i) in the main text. RT-PCR quantification of exon inclusion as PSI values from three independent biological replicates. Statistical significance of differential splicing was determined by the unpaired student’s *t*-test *p*<0.0001 ****; *p*>0.05 ns. **e,** *MTO1* mRNA expression is compromised upon Ku deletion independent of IFN (MAVS KO) or cell cycle states (CDK4/6i) or NHEJ (AID-LIG4).

## List of Extended Data Tables

Extended Data Table 1: A summary of all RNA-seq datasets analyzed in this study.

Extended Data Table 2: Numbers of DSE with six different alternative splicing patterns detected upon depletion of Ku.

Extended Data Table 3: Numbers of DSE cassette exons activated or repressed by Ku, hnRNP C, DHX9, PTBP1, and RBFOX2.

Extended Data Table 4: List of high-confidence Ku-dependent cassette exons with relevant annotations.

Extended Data Table 5: List of genes containing Ku-dependent cassette exons implicated in RNA-binding, DNA repair and mitochondria function.

Extended Data Table 6: List of high-confidence Ku-dependent cassette exons showing differential splicing between brain and non-brain human tissues.

Extended Data Table 7: List of RT-PCR primers used in this study.

