## Extended Data Figs. 1-11 for "Primate-specific adaptation of Ku protects transcriptomic integrity by suppressing Alu-mediated alternative splicing"

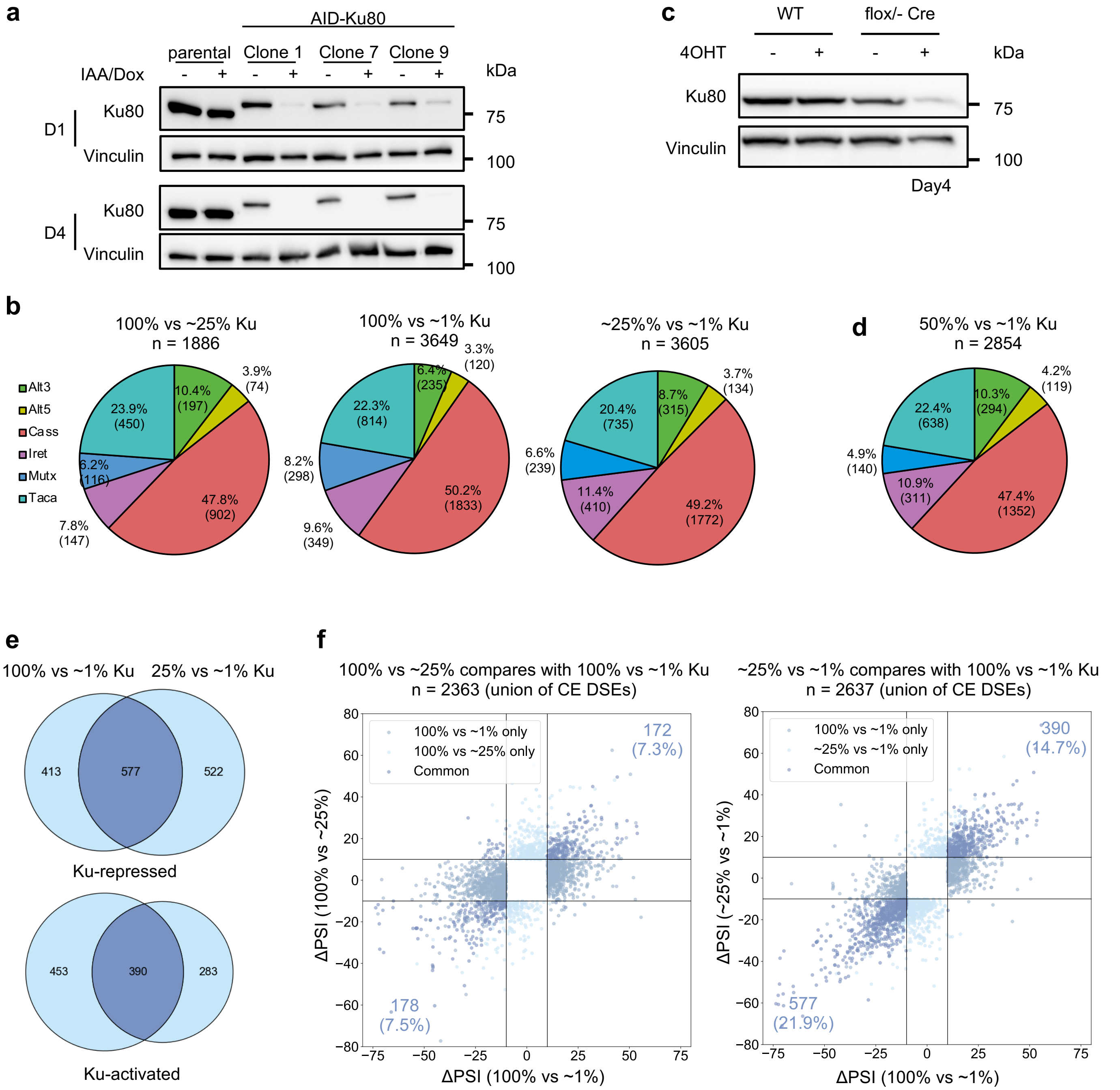


**Extended Data Figure 1**

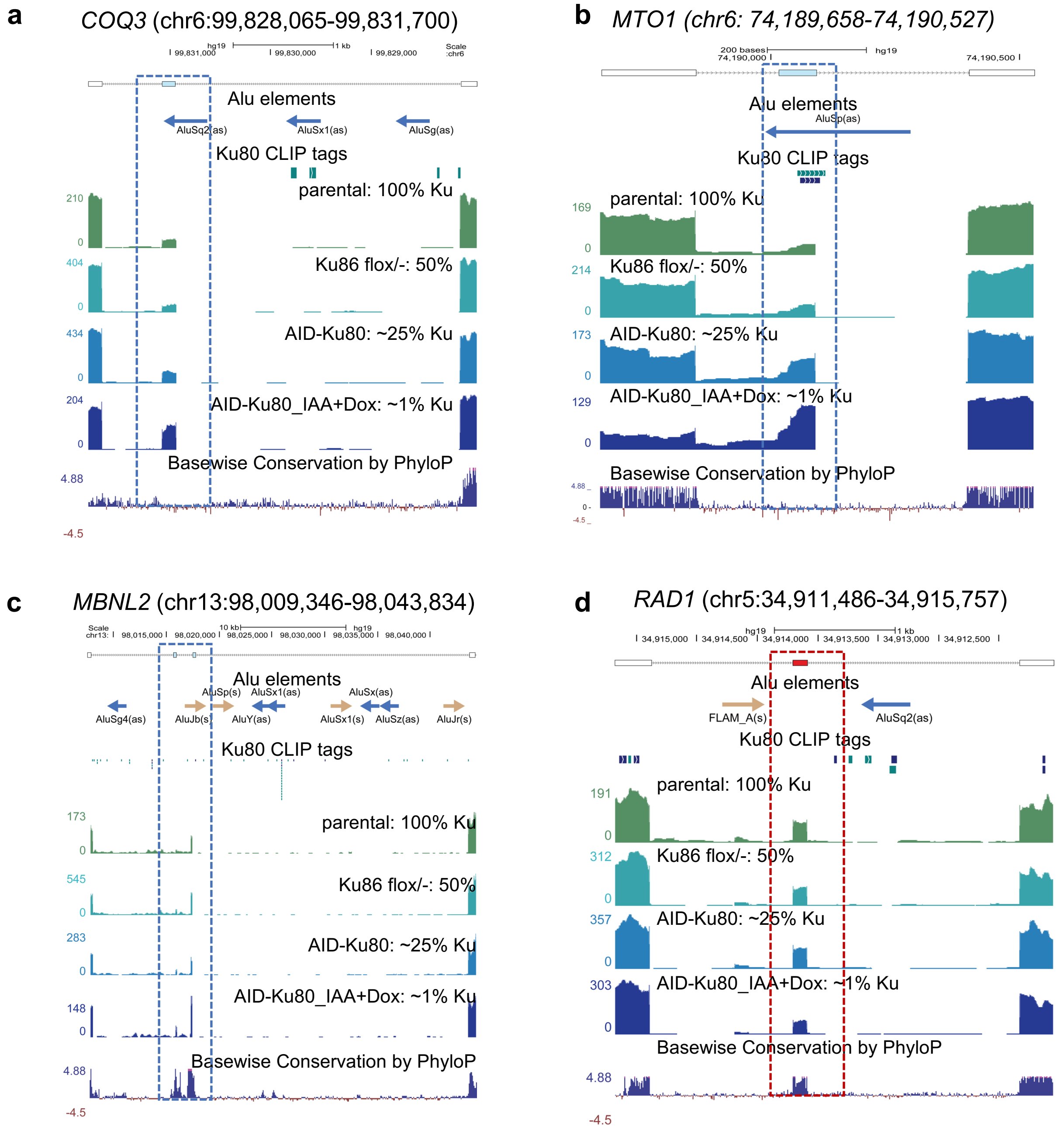


**Extended Data Figure 2**

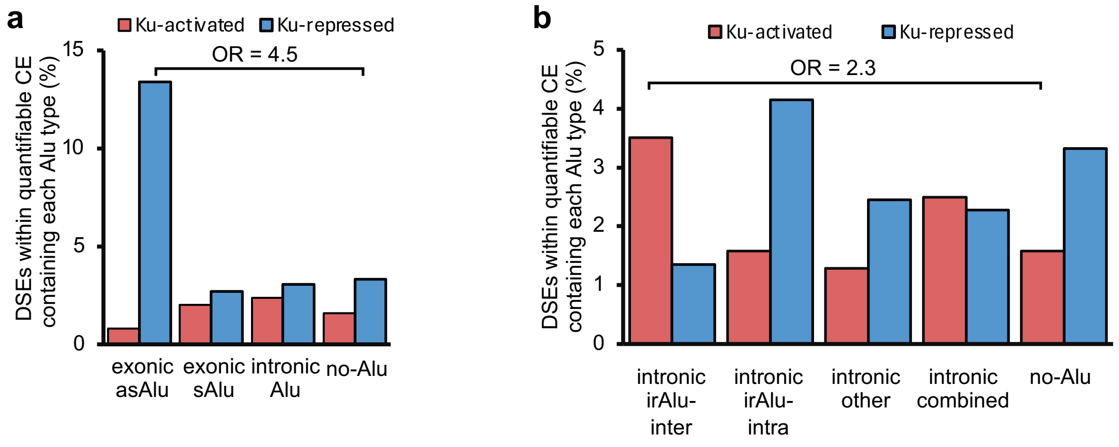


**Extended Data Figure 3**

**a**

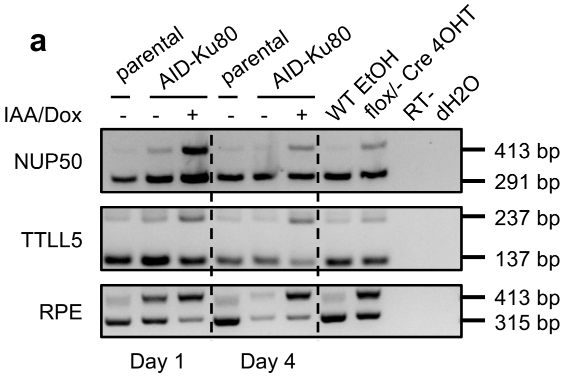


**c**

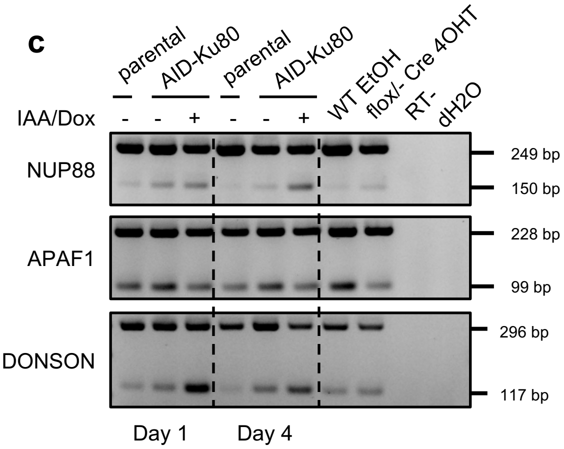


**b**

asAlu (AluSx 288bp) encodes the alternative exon in *EXOSC9*
>hg19_dna range=chr4:122737126-122737413

AG|-: splicing acceptor
-AG|GTAA: splicing donor sites
Underlined: alternative Exon (51nt=>17aa:Isoform 1: 385Q (439aa)=> Isoform 2: QELGFHHVGQTGLEFLTS (456aa)

```
TTTTATTTTT ATTTTTTGGG ATGGAGTCTT GCTCTGTCAC CCAGACTGGA 50
GTGCAGTGGC ACTGCAACCT CCACCTCCAT GGGTTCAAGC GATTCTCATG 100
CCTCAGCCTC GAGTAGCTGG GCCTACAAGC ACATGCCACC ACGCCTGGCT 150
AATTTTTCTA TTATTTTTAA TAGAGCTGGG GTTTCACCAT GTTGGCCAGA 200
CTGGACTCGA GTTCCTGACC TCAGGTAATC CACCTGCCTC AGCCTACCAA 250
AATGTGGGTA TTAAAAGTGT GAGCCACCAC GCCCAGCC
```

**Extended Data Figure 4**

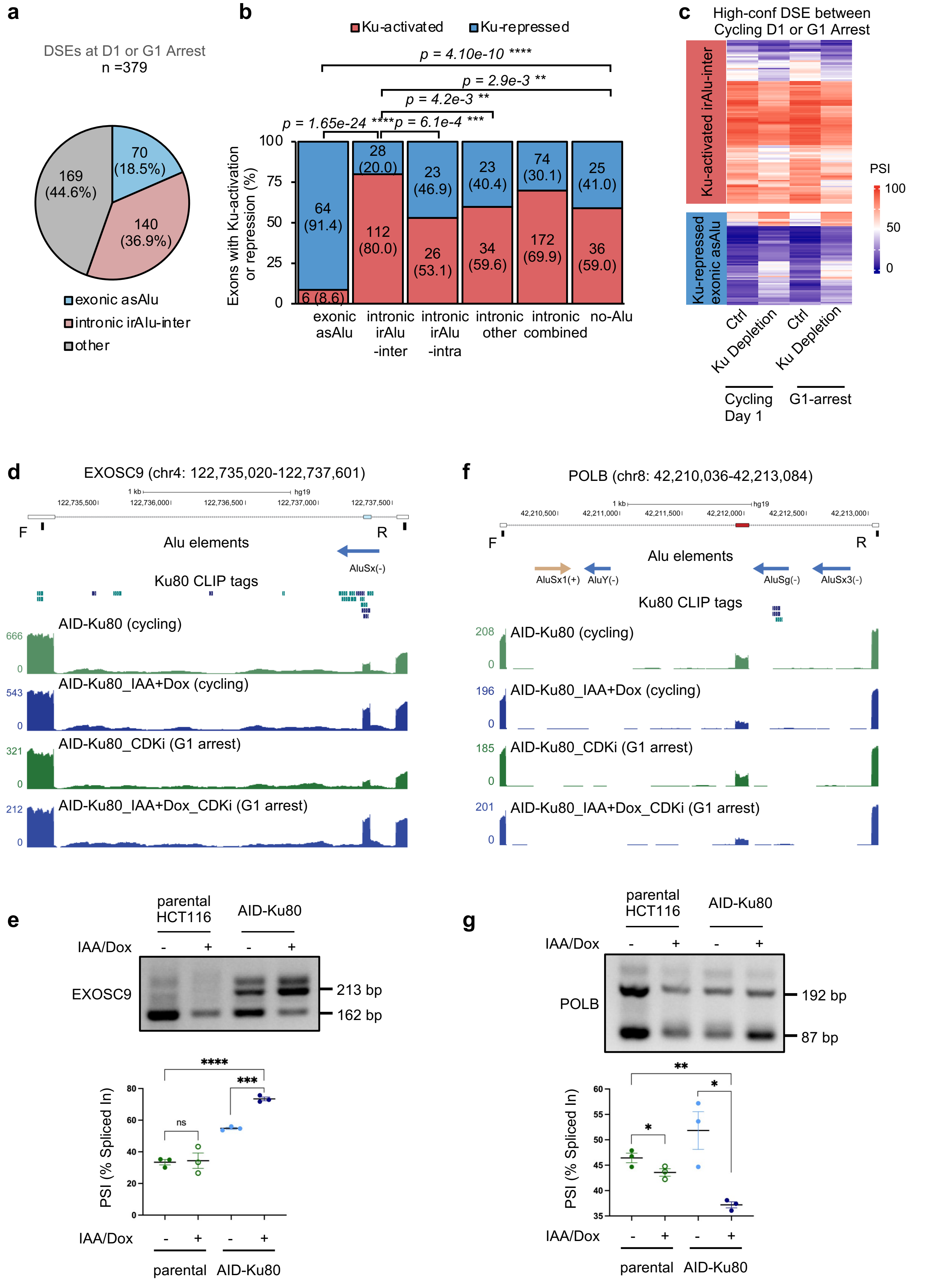


**Extended Data Figure 5**

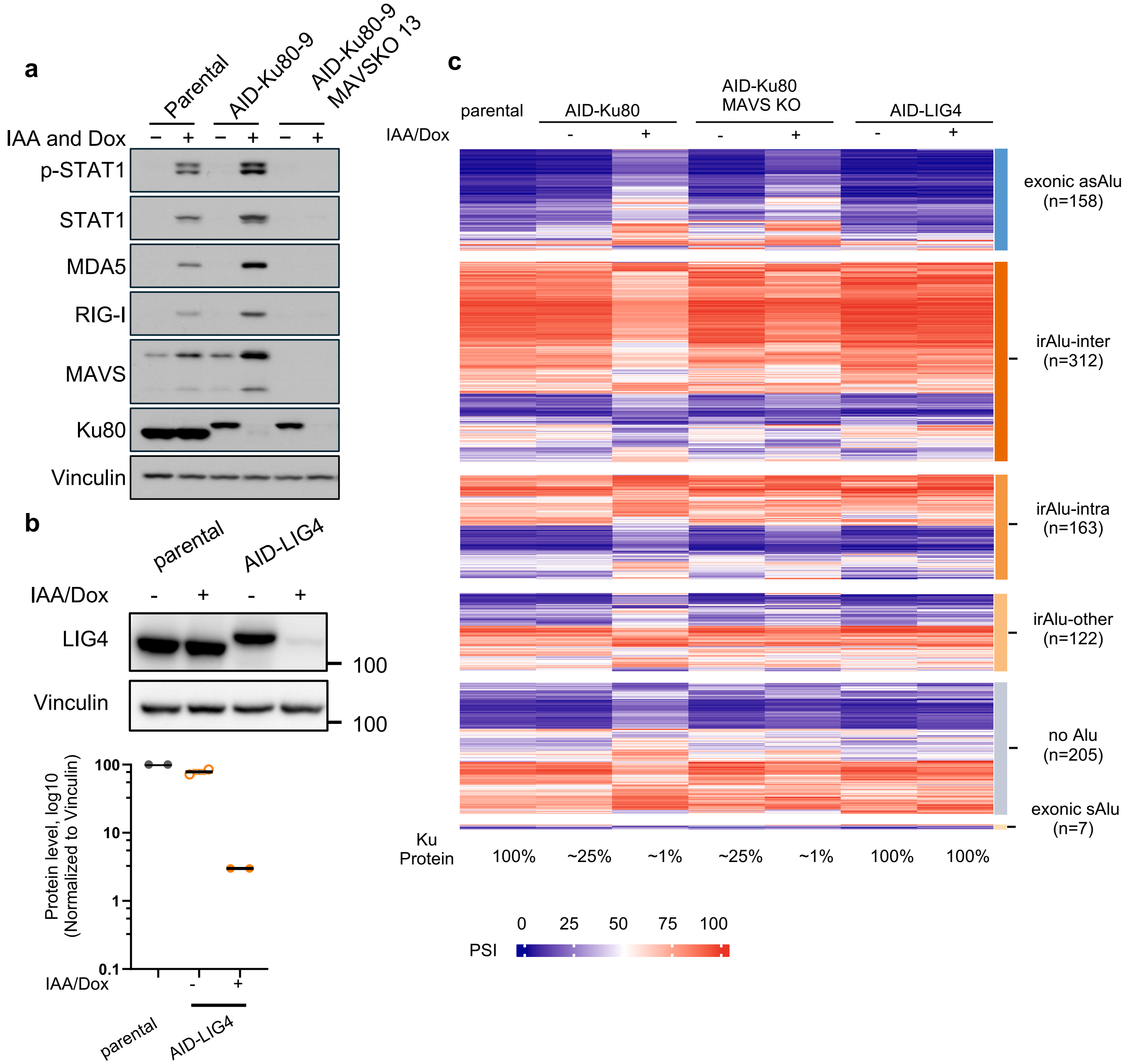


**Extended Data Figure 6**

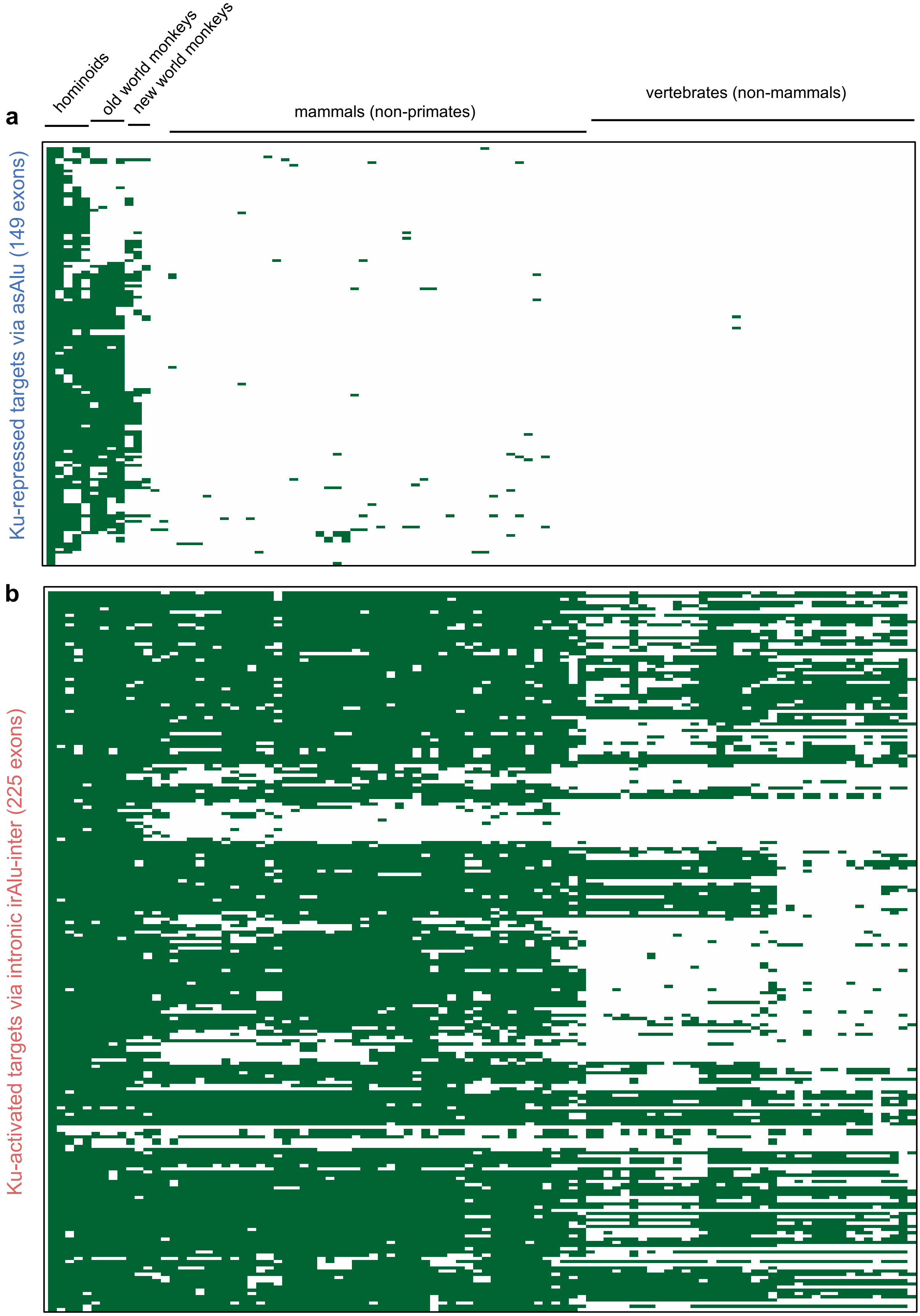


**Extended Data Figure 7**

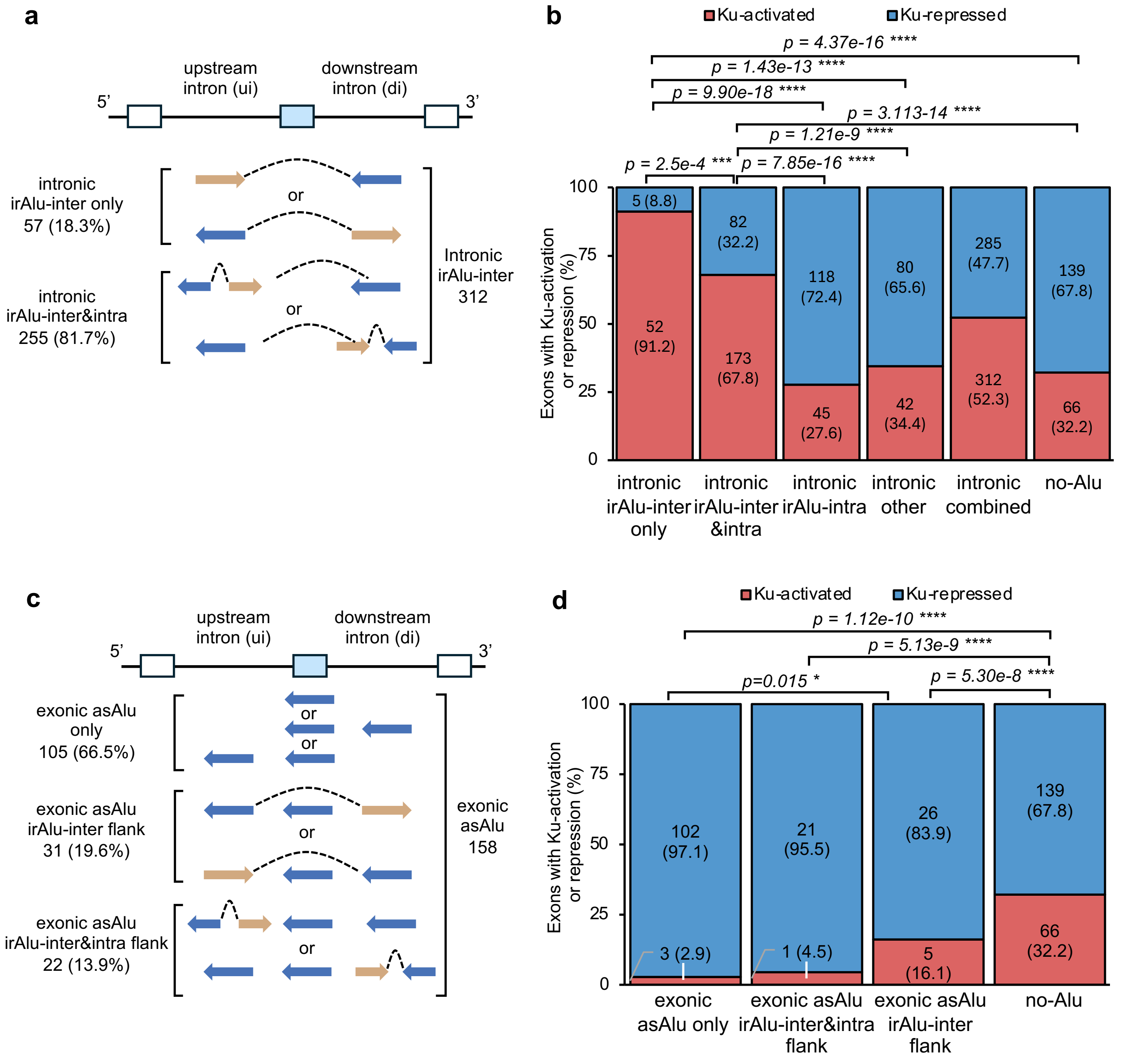


**Extended Data Figure 8**

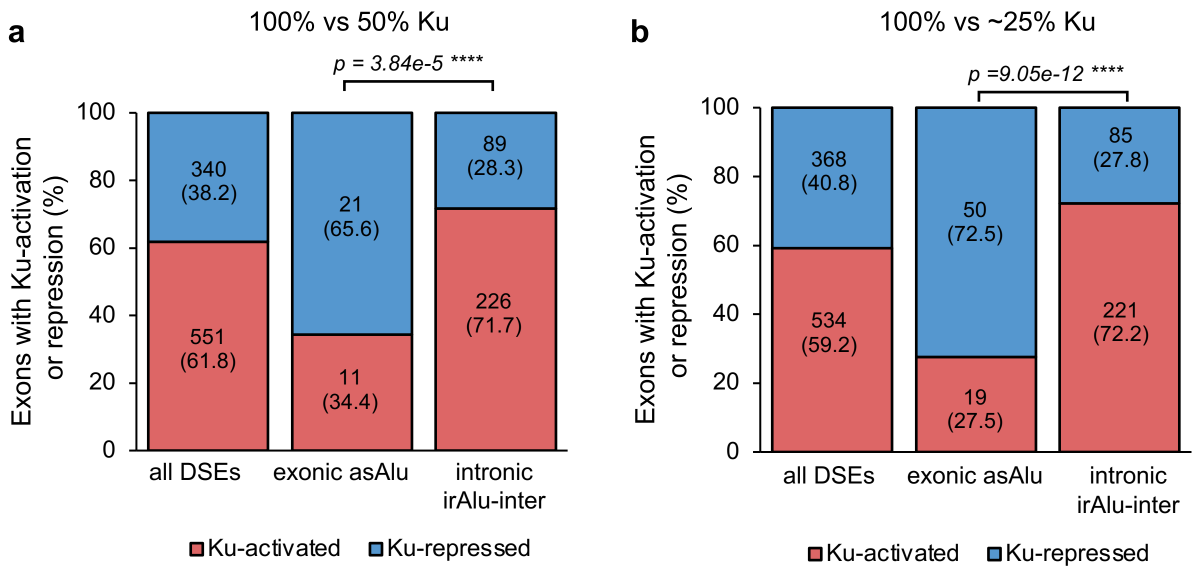


**Extended Data Figure 9**

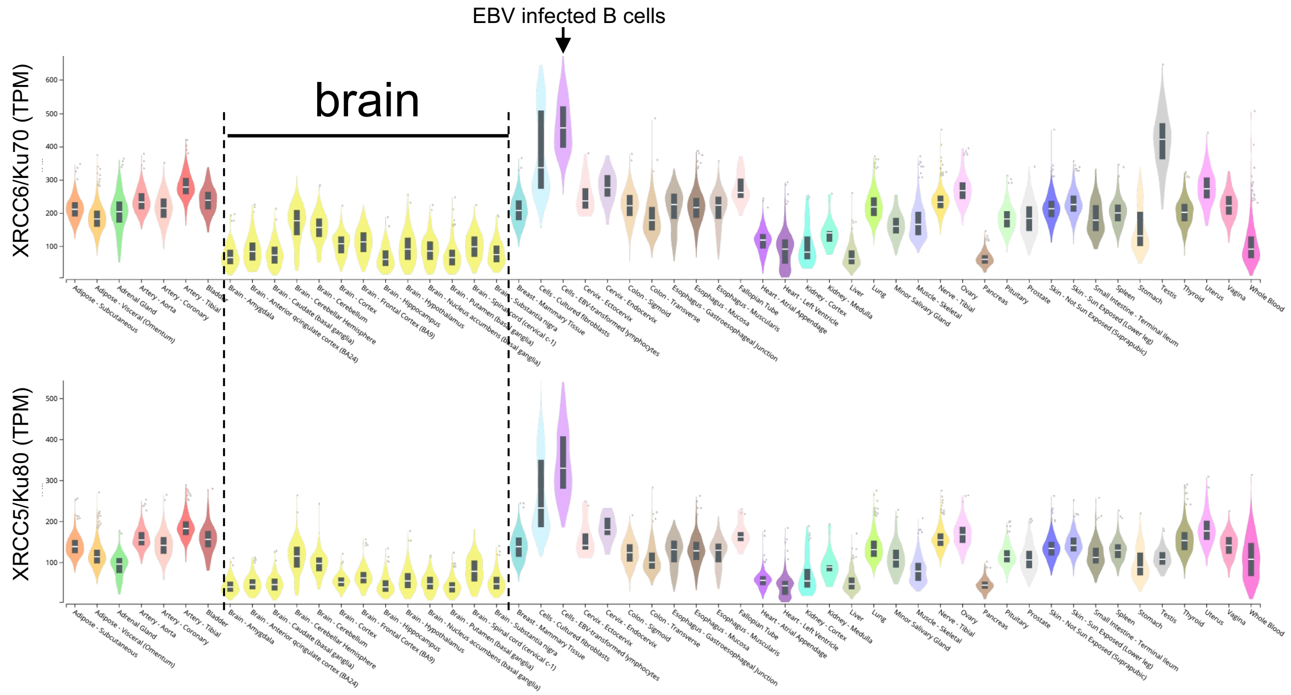


**Extended Data Figure 10**

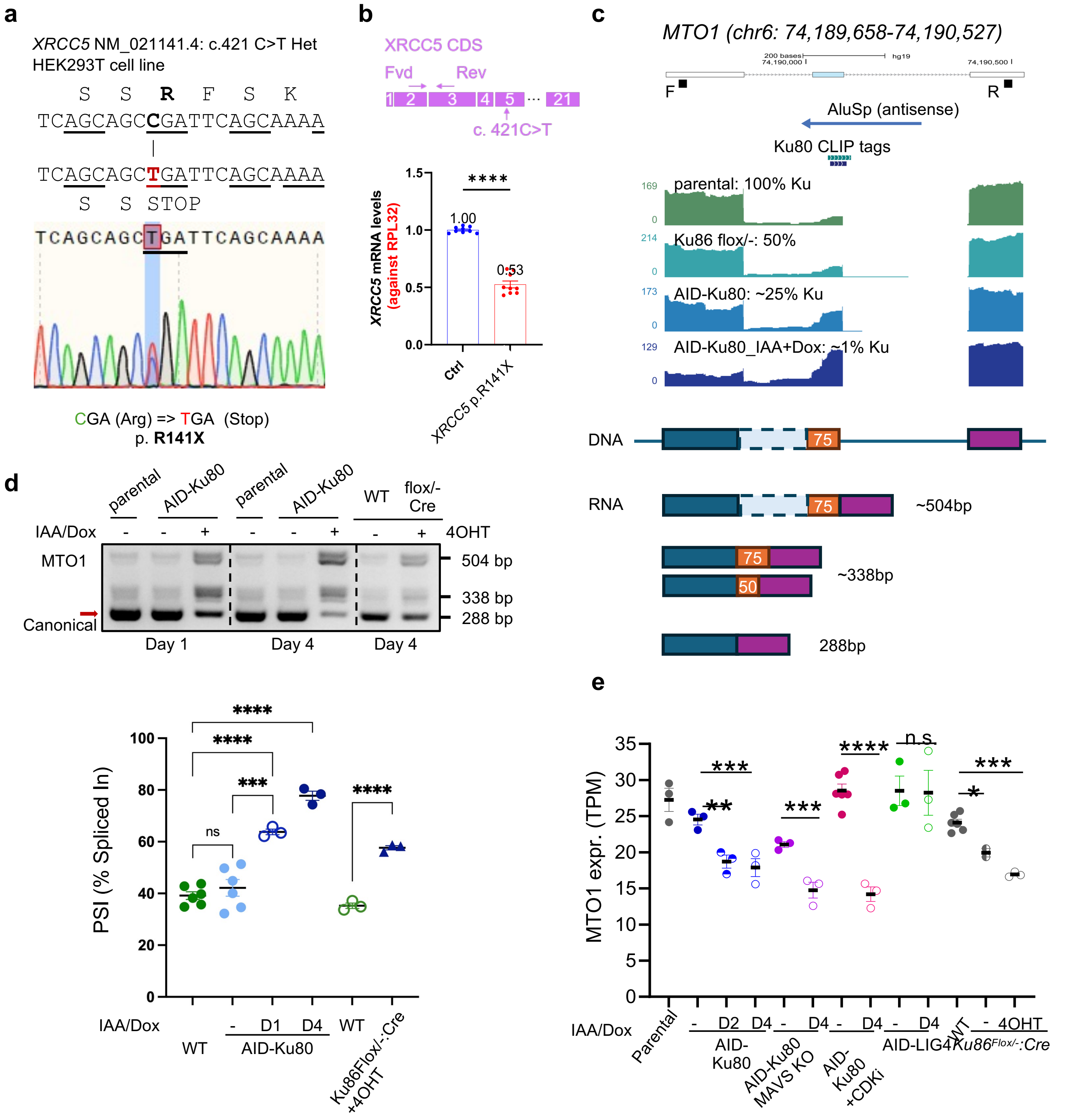


**Extended Data Figure 11**
